# Dryland soil rewetting induces strong VOC emissions with potential to form ozone and aerosols

**DOI:** 10.64898/2026.01.26.701704

**Authors:** Andrea Ghirardo, Benjamin W. Poodiack, Hagar Siebner, Matan K. Jaffe, Baris Weber, Jörg-Peter Schnitzler, Michael Bonkowski, Osnat Gillor, Alex B. Guenther

## Abstract

Biogenic volatile organic compounds (VOCs) significantly influence atmospheric chemistry, yet the importance of microbial VOC emissions remains understudied. We investigate petrichor VOC emissions, the characteristic scent following rainfall after prolonged drought, in an aridity gradient across Israel, by simulating soil rewetting events. Rewetting triggered strong VOC fluxes (1-3.5 nmol m^−2^ s^−1^ ground area) dominated by sesquiterpenes and benzenoids, with emission patterns linked to climatic regions, soil aridity, and microbial community composition. Petrichor consisted of a complex bouquet of 58 VOCs, and the initial VOC burst resembled the CO2 pulse of the Birch effect. Petrichor emissions showed ozone and secondary organic aerosol formation potentials comparable in magnitude to those estimated for anthropogenic VOC sources in Israel. Despite compositional differences, emissions were of similar magnitudes across the aridity gradient. Given that drylands cover nearly half of Earth’s terrestrial surface and are expanding, these episodic microbial VOC emissions may represent a significant, previously overlooked source of reactive carbon with potential implications for regional and global atmospheric chemistry.

## Introduction

Biogenic volatile organic compounds (VOCs) are reactive molecules that play a central role in atmospheric chemistry^1,2^. In the atmosphere, VOCs react with hydroxyl radicals (OH), contribute to ozone (O3) formation, and lead to the production of secondary organic aerosol (SOA), which acts as cloud condensation nuclei. These processes modulate the atmospheric oxidative capacity and the radiative forcing, thereby influencing air quality, cloud formation, and climate^3,4^.

Globally, vegetation is the dominant source of terrestrial biogenic VOCs due to its large biomass and the widespread high terpenoid emissions. Consequently, global VOC emission inventories [e.g., MEGAN^5^], rely primarily on vegetation type and cover. However, these models omit VOC emissions from soils, which are increasingly recognized as a significant yet undervalued source of biogenic VOC. For instance, studies in Amazonia and rainforest mesocosms have reported strong sesquiterpene emissions and other isoprenoids from soils following rewetting^6,7^, implicating soil microbial activity as a potential driver of atmospheric reactivity. These recent findings demonstrate that rainforest soils, particularly during rewetting episodes, can contribute substantially to VOC fluxes, likely via microbial processes. However, the magnitude and importance of soil VOC emissions from other ecosystems remain largely unknown.

Drylands, which include arid, semi-arid, and dry sub-humid regions, cover nearly half of Earth’s land surface (∼66.7 Mkm^2^)^8,9^ and are expanding under climate change^10–12^. Although dryland soils often support sparse vegetation, they host abundant, metabolically versatile microbial communities^13,14^, making them an underexplored source of biogenic VOCs with unknown climate consequences. Water availability strongly structures microbial community composition and function along aridity gradients^13^. In hyper-arid regions, microbes persist via dormancy or hygroscopic water uptake^15^. In contrast, in less arid regions, episodic hydration events rapidly activate microbial metabolism^16^, and cooperative interactions further enhance survival in extremely dry soils^17^.

Biological activity in drylands is tightly coupled with wet-dry cycles. In the Negev Desert, biocrust-dominated communities exhibit rapid resuscitation within minutes after rainfall^8^, and soil rewetting after drought broadly reactivates microbial metabolism, shaping carbon and nitrogen fluxes^18^. A classic example is the Birch effect, a large CO2 pulse following rewetting that can account for a significant portion of the annual carbon budget^19^. Rewetting may also generate VOC bursts, along with microbial reactivation. Upon rewetting after prolonged drought, dry soils are known to release a distinctive scent, namely ‘petrichor’ (from Greek *petra,* meaning stone, and *ichor*, the fluid of the gods), a VOC blend typically associated with geosmin and 2-methylisoborneol (2-MIB) produced by *Actinomycetota* and filamentous *Cyanobacteriota*^20^. While petrichor is familiar to humans, its detailed chemical composition and the extent to which climate- and microbiome-driven adaptations shape emission rates across drylands remain unknown. Petrichor can thus be viewed as a biogeochemical pulse: a short-lived but intense VOC release triggered by soil rewetting, analogously to the CO2 Birch effect, and potentially reflecting both desorption of accumulated compounds and *de novo* biosynthesis. Regardless of whether it is pre-accumulated or newly synthesized, the rapid transfer of petrichor VOCs to the atmosphere, given the vast extent of drylands and the high reactivity of biogenic VOCs, may exert significant impacts on atmospheric chemistry despite its transient nature, underscoring the need to quantify how rewetting pulses influence VOC emissions and carbon fluxes in drylands. Pulse-driven dynamics are a defining feature of dryland ecosystems, where episodic rainfall events control microbial activity, carbon turnover, and trace gas fluxes, often accounting for a substantial fraction of annual biogeochemical exchange^21^.

Here, we investigate soil VOC emissions and carbon fluxes across a steep aridity gradient in Israel, spanning sub-humid (SH) shrubland outside the Negev Desert and semi-arid (SA), arid (A), and hyper-arid (HA) regions within the Negev Desert, which differ in climate, soil properties, and microbial community composition (Fig. 1, Fig. S1-S2, Data S1, Tables S1). Using controlled rewetting experiments that simulate two consecutive rainfall events following contrasting drought histories, we quantify the chemical composition and temporal dynamics of petrichor alongside net ecosystem CO2 exchange under standardized conditions (Methods S1-S2, Fig. S3-S6, Table S2). By upscaling VOC fluxes, this study demonstrates that dryland soils are a significant episodic source of reactive VOCs following rewetting, and that these emissions are shaped by aridity-driven microbial community adaptation and potentially contribute to regional atmospheric chemistry.

**Figure 1.**
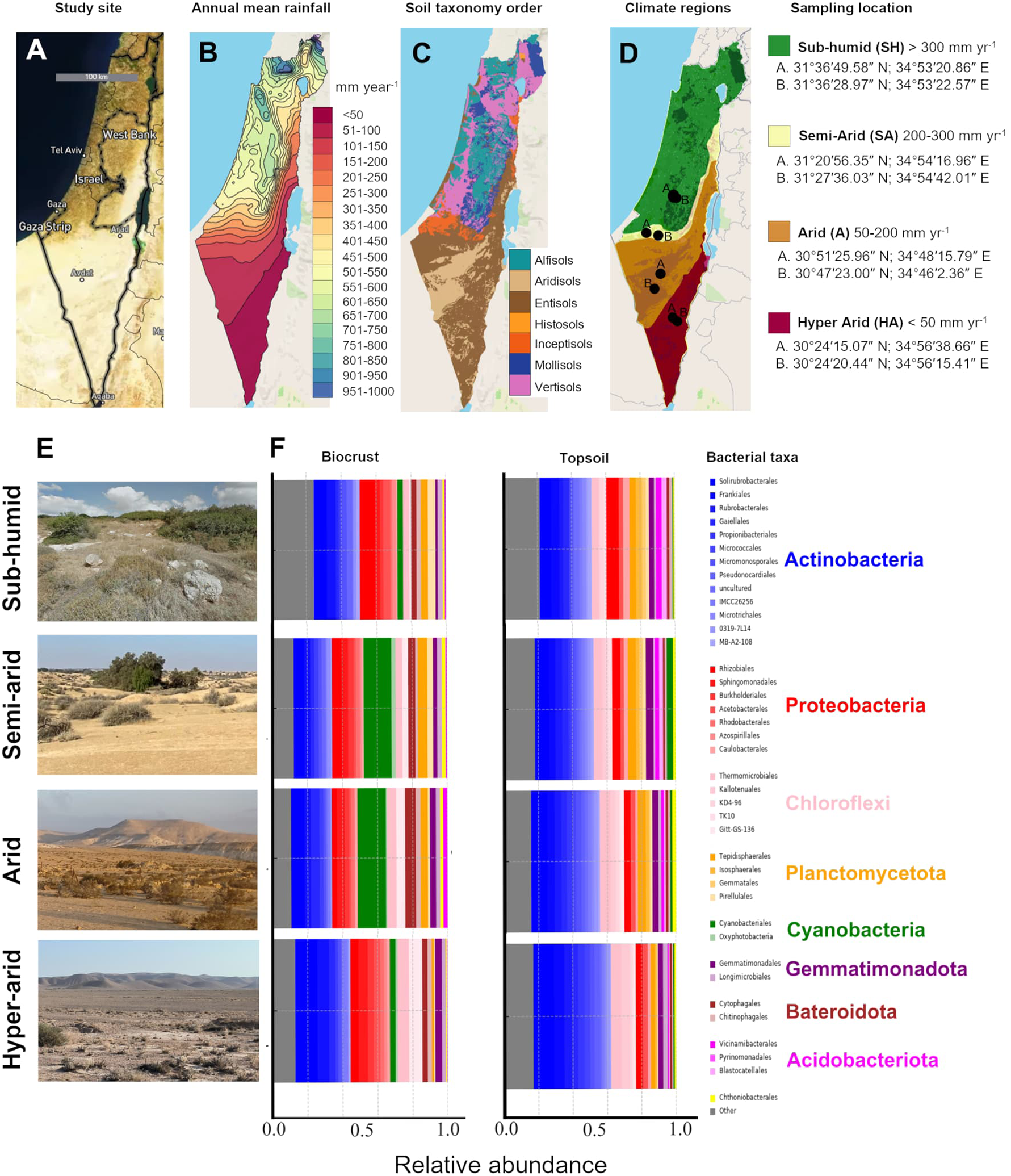
Study sites along the Israel aridity gradient showing climate and soil characteristics, and microbial compositions. (**A**) Satellite image with country administrative borders (black lines). (**B**) Annual mean rainfall across the four sampled climatic regions: subhumid, semi-arid, arid, and hyper-arid. (**C**) Soil taxonomy orders according to USDA classification. (**D**) Overlay of soil taxonomy orders and mean annual precipitation with the geolocation of sampling sites, two spatially separated sites (‘A’ and ‘B’) within each climatic region. (**E**) Photographs of representative collection sites from the sub-humid, semi-arid, arid, and hyper-arid regions. (**F**) Microbial compositions of site-representative sampled soil biocrust and topsoil based on Amplicon sequences of the 16S rRNA genes. Additional details on precipitation, soil moisture and temperature regimes, and soil taxonomy are provided in Fig. S1.

## Results

### Soil bacterial community composition shifts across the aridity gradient

Bacterial community composition in biocrusts and topsoil differed significantly across the aridity gradient (Fig. 1, Table S3-7; *p* = 0.001), with higher within-region variability in arid zones (PERMDISP, F = 11.37, *p* = 0.001). Across both biocrust and topsoil, *Actinomycetota* affiliated orders dominated (56-77%) and increased in relative abundance from sub-humid (SH) toward arid (A) and hyper-arid (HA) regions (Table S3). In biocrusts from semi-arid (SA) and arid regions, *Cyanobacteriales* peaked at 17-18%. Additional soil and region-specific patterns were driven by *Proteobacteria* (14-22% in biocrust), *Chloroflexota* (9-14% in topsoil), and *Bacteroidota* and *Verrucomicrobiota* orders as minor contributors. In biocrusts, multiple *Actinomycetota*, *Cyanobacteriales*, and several *Proteobacteria*, *Planctomycetota*, and *Verrucomicrobiota* orders varied significantly among climatic regions (Table S3-7; FDR-corrected *q*-value < 0.05), whereas topsoil exhibited weaker but overlapping trends. Within each climatic region, many dominant orders differed significantly between biocrust and topsoil, particularly in sub-humid and arid regions, indicating habitat-specific selection between surface biocrust and the underlying soil driven by the aridity gradient (Table S6-7).

### Petrichor composition and emissions depend on aridity gradient and the succession of rewetting events

We quantified the fluxes of VOCs from soil cores collected along a climatic aridity gradient following the two rewetting episodes separated by a month of gradual desiccation, using simulated rewetting experiments and cuvette-based gas-exchange measurements. Rewetting of the soil after 3 months of field drought elicited strong VOC emissions within the first day after the first rewetting, followed by changes in composition and decreased intensity upon the second rewetting (Fig. 2). Petrichor emissions from soils across the climatic regions differed significantly in both intensity and chemical composition during the first and second rewetting episodes, although emissions after the second rewetting were more similar between arid and 3 hyper-arid regions (Fig. 2B-C). No significant differences were detected between different location sites within each climatic region (Fig. 2A), indicating that emission patterns reflect regional climatic adaptation rather than local variability. Across all soils, we detected significant emissions of 58 VOCs, regardless of soil origin. Several benzenoid compounds correlated with immediate emissions and CO2 flux after the first rewetting, whereas several terpenoids were strongly associated with the second rewetting episodes (Fig. 2E-F, Fig. S7, Table S9). Geosmin, a characteristic volatile often identified with petrichor, was also detected in the blend of volatiles after rewetting (Fig S8). However, its emission rate was low (<1 pmol m^−2^ s^−1^) compared to total terpenes (210-640 pmol m^−2^ s^−1^), benzenoids (187-634 pmol m^−2^ s^−1^), and other VOCs (1,050-3,890 pmol m^−2^ s^−1^) mainly composed of the carbonyl compounds acetaldehyde, acetone, and acetic acid, emitted by all soil samples (Fig. 2 G-I). No significant emissions of 2-MIB were detected.

**Figure 2.**
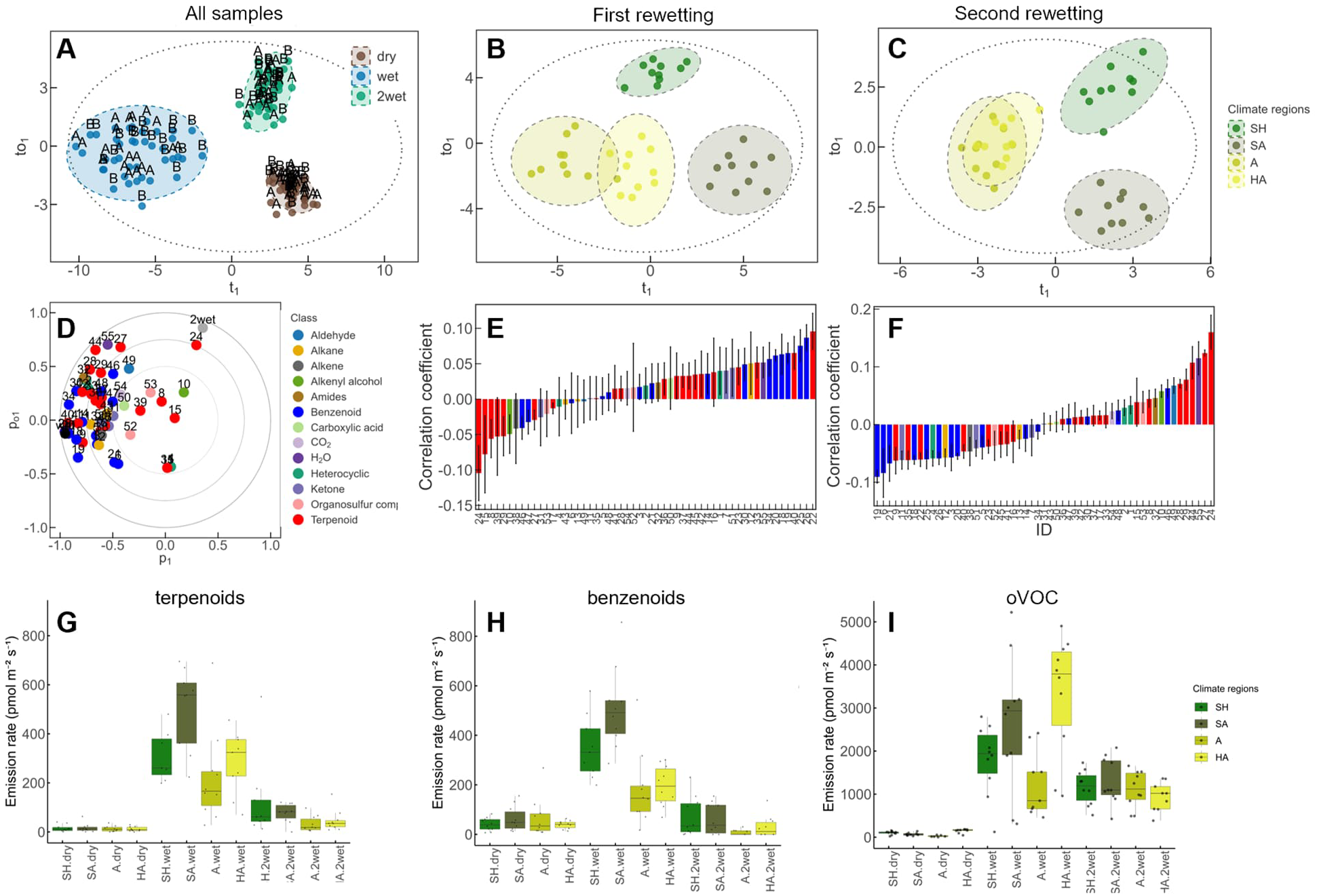
Changes of VOCs emissions upon two consecutive rewetting events. Orthogonal partial least squares (OPLS) analysis of VOC emission rates from petrichor in sub-humid (SH), semi-arid (SA), arid (A), and hyper-arid (HA) climatic regions, under dry conditions and following two consecutive rewetting events. (**A**) Score plot for all samples combined; (**B**) after the first rewetting; (**C**) after the second rewetting; (**D**) normalized and correlation-scaled loading plot for all samples. (**E-F**) Correlation coefficients of variables explaining the climate-region separation for: (E) first rewetting; (F) second rewetting (ID = VOC name, given in Table S9). Box plot of VOC emission rates of total (**G**) terpenoids; (**H**) benzenoids; (**I**) oxygenated small VOCs (oVOCs) that include acetaldehyde, acetone, and acetic acid (n=10). Each point represents 17 h integrated flux of an individual replicate (soil core) collected from two distal sites (2-10 km apart; ‘A’ and ‘B’) within the same climate region. Ellipses indicate 95% confidence intervals around group centroids; the large outer ellipse represents Hotelling’s *T*^2^ (95%) confidence region for all samples. All OPLS models were statistically significant (CV-ANOVA: A, p < 1×10^−10^; B, p = 1×10^−8^; C, p = 1×10^−5^). Model fitness: A, R^2^X(cum) = 0.59, R^2^Y(cum) = 1, Q^2^Y(cum) = 0.87; B, R^2^X(cum) = 0.63, R^2^Y(cum) = 1, Q^2^Y(cum) = 0.76; C, R^2^X(cum) = 0.68, R^2^Y(cum) = 1, Q^2^Y(cum) = 0.53. VOCs were collected the day before and immediately after each rewetting event.

The aridity gradient significantly (*p* < 0.001, CV-ANOVA) impacted petrichor emissions during both the first and second rewetting events (Fig. 3). Terpenoid emissions were characteristic of the wetter sub-humid region, particularly monoterpenoids such as eucalyptol and camphor (Fig. 3, S9-10, Table S9). Benzenoids (e.g., xylenes, indane) and the sesquiterpene longicyclene were associated with semi-arid, whereas both benzenoids and the monoterpenoids 2-methyl-2-bornene and (*E*)-p-menthan-1-ol correlated with the arid region. Emissions of isolongifolene, acetaldehyde, and alkanes (e.g., decane) were positively correlated to the hyper-arid region. Following the second rewetting, the aridity gradient was also associated with shifts in mono- and sesquiterpenoid emission profiles (Fig. 3F, 3H). Overall, C, N and P, Fe and Mn availability were higher in the wetter sub-humid and semi-arid regions, whereas soluble ions (e.g., NO3^−^, Cl^−^, Na^+^) were significantly higher in the drier arid and hyper-arid regions (Fig. S2).

**Figure 3.**
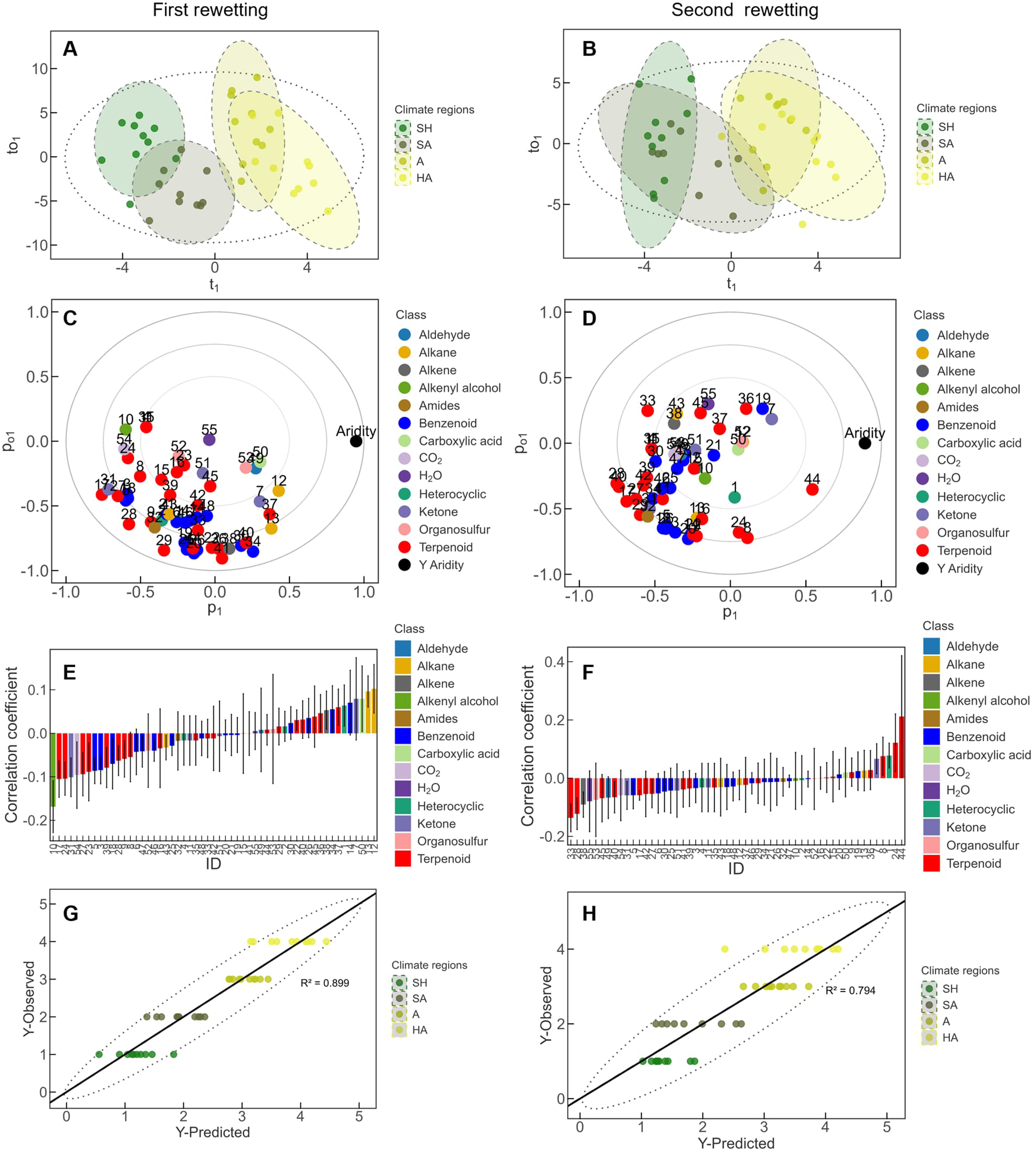
Differences of VOC emissions along an aridity gradient. OPLS analysis of VOC fluxes showing dependencies of petrichor emissions and the aridity gradient after the first (left panels) and second (right panels) rewetting events. Climatic regions: sub-humid (SH), semi-arid (SA), arid (A), and hyper-arid (HA). (**A-B**) Score plot; (**C-D**) normalized and correlation-scaled loading plot. (**E-F**) Correlation coefficients of variables explaining the aridity gradient. (**G-H**) Predicted *vs* Observed Y-values of the aridity gradient showing significant linear impact of aridity on petrichor emissions (first rewetting: p < 1×10^−9^; second rewetting p = 1.13×10^−4^, CV-ANOVA). Root Mean Square Error of Estimation (RMSEE = 0.375) and RMSE Cross-validated (RMSEcv = 0.537). Ellipses indicate 95% confidence intervals around centroids; the large outer ellipse in C-D represents Hotelling’s *T*^2^ (95%) confidence region. Model fitness: A, R^2^X(cum) = 0.53, R²Y(cum) = 1, Q^2^Y(cum) = 0.90; B, R^2^X(cum) = 0.32, R²Y(cum) = 1, Q^2^Y(cum) = 0.79. Numbers in C-F indicate the VOC ID listed in Table S9.

### Emission dynamics and correlations of VOCs, H2O and CO2 fluxes

Soil rewetting is known to induce sharp CO2 pulses, yet the temporal dynamics of VOC emissions during these events remain unknown. To capture these rapid metabolic and physicochemical shifts, we simultaneously measured real-time fluxes of VOC, H2O and CO2 from the soil cores over four consecutive days after each rewetting (Fig. 4). As expected, rewetting caused a rapid and pronounced pulse of CO2 (the Birch effect). Notably, VOC emissions showed similar temporal dynamics to CO2, with acetaldehyde exhibiting the strongest similarity (Fig. 5).

**Figure 4.**
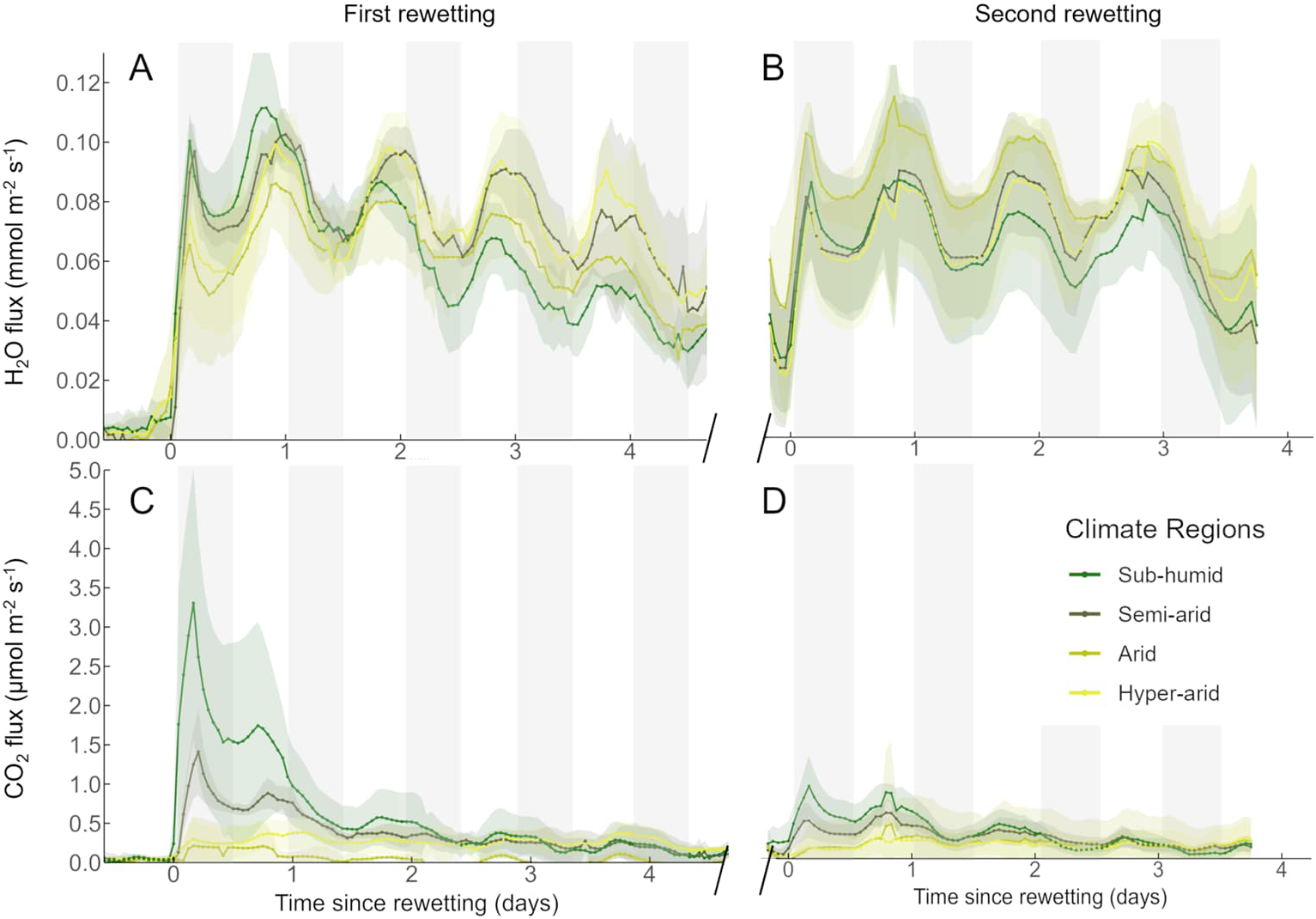
Soil H2O and CO2 flux responses to two rewetting events across an aridity gradient under controlled environmental conditions. Mean fluxes of H2O (**A-B**) and CO2 (**C-D**) measured in soils from the four climatic regions. Left and right panels refer to the first and second rewetting events, respectively. After the first rewetting, the soils were allowed to dry for 31 days at standard conditions (T = 23 ± 0.4 °C; RH 53 ± 5%). Time is expressed relative to the onset of rewetting (day 0), and the day/night cycle is indicated with white/grey bars. Lines represent mean fluxes and shaded areas the 95% confidence intervals (n = 10).

**Figure 5.**
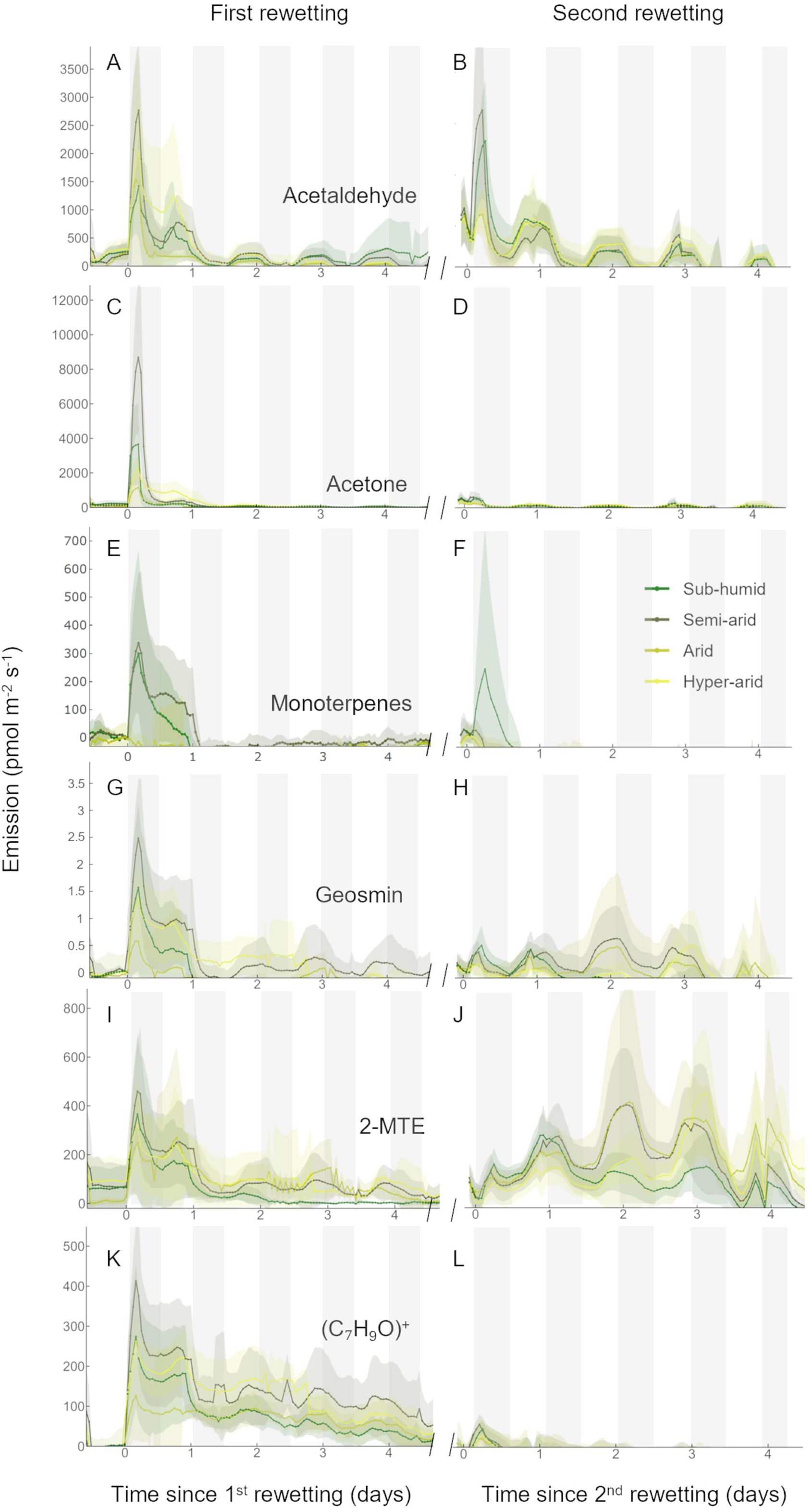
Emission dynamics of VOCs in the petrichor in response to two rewetting events. Time series of VOC emission rates (normalized to ground area) measured by PTR-MS of acetaldehyde (**A-B**), acetone (**C-D**), monoterpenes (**E-F**), geosmin (**G-H**), 2-methylthioethanol (2-MTE) (**I-J**), benzenoid and aromatic alcohols at C7H9O+ (**K-L**). Left and right panels refer to emissions following the first and second rewetting event, respectively, separated by 31 days of desiccation. Soil cores originate from the four climatic regions: sub-humid, semi-arid, arid, hyper-arid. Time is expressed relative to the onset of rewetting (day 0), and the day/night cycle is indicated with white/grey bars. Lines represent mean fluxes and shaded areas the 95% confidence intervals (n = 10).

Water vapor fluxes across the aridity gradient were essentially similar after the first and the second rewetting (Fig. 4A-B), reflecting the similar evaporation process under the controlled conditions and differences in the soil texture (Fig. S2, Table S1). Conversely, fluxes of CO2 were much higher after the first rewetting event following drought and then declined with the second rewetting, suggesting that some labile carbon pools were depleted after a short desiccation-hydration cycle. When comparing the soils, CO2 fluxes were significantly higher from soil collected in regions with higher precipitation (the sub-humid and semi-arid regions), compared to the drier arid and hyper-arid regions (Fig. 4). The daylight cycles affected both the H2O and CO2 fluxes, driven by temperature and relative humidity changes inside the cuvettes [28.4/22.8 °C, 11/65% RH (light/dark), Fig. S4]. Importantly, CO2 fluxes were nearly zero before both rewetting episodes (dry conditions), indicating that the microbial metabolic activity was successively suppressed by desiccation (Fig. 4C, D).

VOC emissions generally peaked within hours of rehydration, exhibiting distinct temporal profiles and intensities across compound classes and between wetting events (Fig. 5). Acetaldehyde emissions were rapidly and strongly induced after the first rewetting (1,000-3,000 pmol m^−2^ s^−1^, peaking at 3-4 hours after rewetting), with comparable peak magnitudes following the second rewetting (Fig. 5A-B). In contrast, the acetone emissions were markedly higher during the first rewetting (up to 8,000 pmol m^−2^ s^−1^, peaking at 3-4 hours after rewetting) but declined significantly during the second event (220-580 pmol m^−2^ s^−1^) (Fig. 5C-D). Larger ketones (C4-C10) followed a similar trend (Fig. S11). Acetic acid mirrored acetaldehyde, decreasing from the first to the second rewetting events, but the reduction was smaller than that of ketones. Terpene emissions were strongly elicited upon rewetting (Fig. S12). Monoterpene emissions were particularly associated with soils from the sub-humid regions (Fig. 5E-F), which contained small particulate of plant materials (Fig. S6). Geosmin emissions from semi-arid region increased gradually over time following an initial transient pulse and remained consistently higher than those from other soil types throughout the measurement period (Fig. 5G-H). In contrast, monoterpenoid emissions were low during the second rewetting event and were primarily associated with sub-humid and semi-arid soils.

The sulfurous volatile compounds dimethyl sulfide (DMS) and 2-methylthioethanol (2-MTE) exhibited contrasting emission patterns across wetting events. A pronounced DMS burst was observed following the first rewetting, particularly in soils from the sub-humid region (Fig. S13). In contrast, 2-MTE emissions increased notably after the second rewetting (Fig. 5I-J), with the highest fluxes observed in soils from the semi-arid and arid climatic regions. Benzenoid and aromatic alcohol compounds, detected at m/z 109 [(C7H9O)⁺], and xylenes (m/z 107), showed a rapid decline over time (Fig. 5K-L; Fig. S13). Although the second rewetting induced a temporary emission, levels dropped quickly and became undetectable within two days.

Diurnal cycles in VOC fluxes were evident for several other compounds (e.g., MT, geosmin, aldehydes), suggesting a strong influence of temperature and light on emission dynamics (Fig. 5, S4, S11-13). The emission dynamics of acetaldehyde, acetone, acetic acids, and 2-MTE were strongly interrelated (clustered heatmap, Fig. S14). In particular, acetaldehyde and acetic acid showed strong correlations (Spearman’s correlation coefficients > 0.91). When comparing individual volatiles measured by GC-MS and PTR-MS, emissions of several terpenoids and benzenoids were also found to correlate (Fig. S15), suggesting a common microbial source or shared biochemical origins.

### Petrichor emissions impact OFP and SOAP

To assess the potential impact of the strong petrichor VOC emissions on atmospheric chemistry, we calculated the ozone forming potential (OFP) and secondary organic aerosol potential (SOAP) expected after the first rewetting event (Fig. 6). While OFP and SOAP provide more meaningful metrics than mass emissions by accounting for VOC reactivity and SOA yields, respectively, actual production rates depend on non-linear chemistry involving NOx, sunlight, and environmental conditions that vary with location and time. Mass emissions, OFP, and SOAP (g m^−2^ day^−1^) were calculated for each compound, summed by chemical category (benzenoids, monoterpenoids, sesquiterpenoids, other), and determined separately for each soil and climatic region. The area-specific emission rates across the aridity gradient were then upscaled to land areas to estimate total petrichor VOC emissions for Israel, yielding an estimate of 400 metric tons (MT) per day. The Petrichor OFP and SOAP estimates were 1120 MT day^−1^ (55 kg km^−2^ day^−1^) for OFP and 27.6 MT day^−1^ (1.36 kg km^−2^ day^−1^) for SOAP.

**Figure 6.**
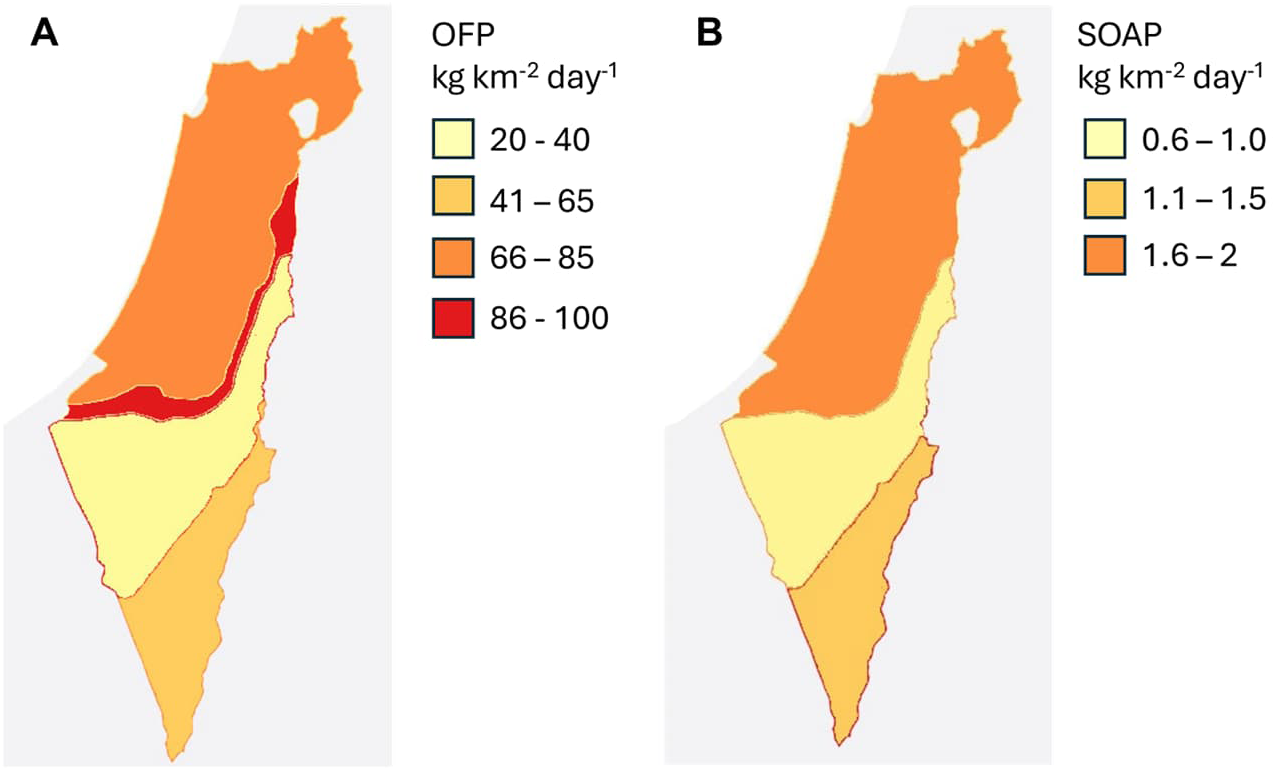
Petrichor-driven O3 and SOA formation potential in Israel. (**A**) OFP, and (**B**) SOAP were calculated from climate-region-specific petrichor VOC fluxes from soils measured the day after rewetting, using maximum incremental reactivity (MIR) and SOAP weighting factors for each compound. The figures show OFP and SOAP formation associated with post-rewetting petrichor emissions across regions.

Although the semi-arid region had >40% higher mass emissions than the sub-humid region, its OFP was only 14% higher and its SOAP similar (−1%) (Fig. S16). The hyper-arid region mass flux was only −11% lower than in sub-humid, while the OFP and SOAP were about 30% lower. In both cases, the differences between the mass flux and the OFP/SOAP results were due to the relatively high contribution of monoterpenes to the sub-humid flux. In contrast, the arid region fluxes were consistently ∼58% lower than in sub-humid region for mass, OFP, and SOAP due to their similar composition.

## Discussion

### Strong VOC fluxes upon rewetting after drought

Rewetting dry soils after a long drought across a steep aridity gradient triggered strong pulses of VOC emissions, analogous to the Birch effect^20,22^. These bursts likely reflect a combination of abiotic desorption processes and biotic responses, including osmotic-shock-induced cell lysis, rapid mineralization of labile organic matter, and the immediate activation of microbial metabolism ^20,23,24^. The rapid microbial resuscitation observed in desert soils^8^, together with the post-rewetting rise in terpenoid and sulfur-containing volatiles, suggests a contribution from *de novo* biosynthesis alongside passive release. In rainforests and Mediterranean soils, rewetting was shown to stimulate soil VOC emissions, consistent with rapid microbial activation following hydration^7,20^. Together, these observations suggest a common mechanism across biomes where the first rewetting event after drought triggers soil VOC emissions.

### Microbial and physicochemical controls on the petrichor emissions

Petrichor composition differed across the aridity gradient, reflecting differences in climatic history, water availability, soil organic matter, and microbial communities (Fig. 1). Yet, the same core of VOCs was detected across climatic regions, albeit in different proportions. Aldehydes, ketones, benzenoids, and terpenoids dominated the first rewetting pulse (Fig. 2, S11), consistent with microbial stress responses, rapid carbon processing, and byproduct release under low-oxygen conditions in rewetted arid topsoil^20,25^. The emission dynamics likely reflect coupled biochemical and physicochemical processes involving metabolites that accumulate during drought and are subsequently released upon soil rewetting, either from damaged microbial cells or from soil pores ^7,24,26^. During desiccation, microbes accumulate osmolytes to maintain cellular integrity, and upon rewetting, the abrupt drop in osmotic pressure promotes cell lysis, osmolyte release, and rapid microbial resuscitation^8,24^. The osmolytes consist of sugars and amino acids, that are rapidly catabolized, producing CO2 and oxygenated VOCs such as acetate, acetone, aldehydes, alcohols, and small organic acids^26,27^. Physicochemical processes also cause VOC emissions^20^. During drought, organics are adsorbed in soil micropores, and when rewetted, changes in gas diffusion, sorption equilibria, and soil polarity promote desorption and volatilization, leading to a rapid VOC pulse^24^. Together, moisture-driven physicochemical release and microbial lysis likely explain the rapid, strong petrichor pulse observed immediately after the rewetting episodes across the aridity gradient, whereas sustained, secondary emissions suggest *de novo* microbial metabolism. In general, although PTR-MS resolved the rapid VOC emission dynamics following rewetting, temporal emissions alone cannot distinguish *de novo* microbial production from physicochemical release, which would require stable isotope labeling studies; nonetheless, compound-specific dynamics, their coupling to CO2 fluxes, and their systematic variation across the aridity gradient support a dominant microbial control on sustained petrichor emissions.

The dynamics of benzenoid emissions likely reflect a combination of rapid desorption from soil surfaces and microbial production linked to shikimate pathway. Some benzenoids (e.g., ethenylbenzene) are known to originate from microbial phenylpropanoid metabolism, especially in Proteobacteria and fungi^28,29^. However, hydrophobic aromatics such as xylenes, ethyl- and polymethyl-benzenes and indanes may partly derive from anthropogenic combustion residues that are transported over long distances on black-carbon particles, which are abundant in dryland soils and exhibit a strong affinity for aromatic hydrocarbons^30,31^. Rewetting alters sorption equilibria and pore accessibility, enabling desorption, as discussed above. This likely explains the strong attenuation of these emissions upon a second rewetting under standardized conditions.

Terpenoids also peaked immediately after rewetting from soils of the sub-humid regions, but they also increased over time. Terpenoid biosynthetic genes are widespread in soil bacteria and fungi, acting as cell membrane stabilizers, oxidative protectants, and infochemicals mediating microbial communication and competition^15,32^. Their delayed appearance suggests linkage to microbial growth phases and de novo processes. The initial emission pulse likely reflects the physical release of pre-existing terpenoids associated with the producer cell membranes and/or stored pools that accumulated during desiccation, while the subsequent gradual increase suggests de novo biosynthesis during the microbial regrowth phase. Rewetting stimulates microbial resuscitation^33^, which can lead to renewed metabolic activity in terpene-producing genera such as *Streptomyces* (*Actinomycetota*), *Collimonas* (Proteobacteria), and filamentous fungi^32^. As these organisms shift from maintenance to growth, carbon flow through terpene pathways may increase. The delayed increase in terpenoid emissions, therefore, may reflect microbial succession and metabolic recovery processes rather than purely abiotic desorption. This interpretation is consistent with the time scales of rapid resuscitation and secondary metabolism following rewetting^8,13^, and could be tested again using stable isotope approaches. While we cannot fully disentangle microbial biosynthesis from physicochemical release, the compound specificity, temporal evolution, and coupling to CO2 fluxes and microbial community structure indicate a dominant microbial control on sustained emissions.

Microbial sulfur volatiles such as DMS and 2-MTE are important mediators of plant-microbe and microbe-microbe interactions^32^. Interestingly, we observed contrasting dynamics of sulfur VOCs: DMS peaked immediately after the first rewetting, possibly reflecting stress-related release, while 2-MTE dominated after the second wetting (Fig. 5, S13), suggesting slower microbial sulfur metabolism. Similar sulfur pulses have been observed in rainforest rewetting experiments^7^. Metabolic production of 2-MTE is intriguing, as it is a product of sulfur recycling by soil bacteria that possess a methylthio-alkane reductase^34^. Some bacteria can use 2-MTE to produce methionine and S-adenosylmethionine under limited sulfur conditions and in anoxic environments^34^, a strategy to scavenge sulfur in anaerobic soil niches. Such anoxic microsites have been documented in rewetted biocrusts and desert soils^35^. Yet, linking these dynamic sulfur signals to functional expression will require targeted omics and mechanistic studies.

### Microbial community and aridity gradient effects

Soil microbial community composition shifted strongly across the aridity gradient^13^ (Fig. 1). Sub-humid and semi-arid soils showed higher CO2 fluxes, reflecting greater total organic-carbon content, whereas arid and hyper-arid soils emitted proportionally more geosmin and sulfur volatiles, consistent with the dominance of stress-tolerant, trace gas-oxidizing *Actinomyceota* taxa^13^. These trends align with metagenomic evidence that desert communities rely increasingly on chemosynthetic and hydrogenotrophic pathways under extreme aridity ^35^.

As the aridity gradient also implies shifts in nutrient availability (C, N, and P) and salinity (Suppl Fig. S2), these chemical differences likely further shaped the volatilome. Nutrient-richer sub-humid soils appear to promote *de novo* microbial VOC biosynthesis during post-wetting regrowth, whereas nutrient-poor, saline arid and hyper-arid soils showed VOC signatures consistent with osmotic-shock responses, characterized by early emission pulses dominated by low-molecular-weight carbonyls, alkanes, and benzenoid compounds^20,24^.

We hypothesize that the VOC emissions help shape the microbial community structure, as many microbial volatiles function as allelochemicals that inhibit competitors or modulate growth and survival^36^. Microbial volatiles can alter gene expression, modulate physiology, and biofilm formation in VOC-exposed bacteria^37^. The microbial communities resuscitate within minutes of hydration in these drylands, allocating much of their early activity to cellular repair, energy generation, and competitive interactions^8,36^. The climate-specific and temporal shifts in VOC composition observed here may serve as fingerprints of microbial community structure, physiological activity, stress responses, and interspecific communication, yet their ecological roles in desert soils remain to be discovered. These volatile signals likely act at short and large spatial scales, with unclear broader ecological consequences ^38–40^.

### Unveiling the potential atmospheric impact of soil VOC emissions from dryland

Petrichor VOCs are highly reactive compounds in the atmosphere. Early pulses of benzenoids and alcohols may act as sinks for hydroxyl radicals, while later emissions of terpenoids serve as precursors for SOA and contribute to OH recycling^1,2^. DMS is a key biogenic sulfur gas that affects aerosol and cloud formation^41^. Thus, the chemical composition of desert VOC fluxes is as crucial as their emission magnitude.

Quantitatively, using the daily mean averages, we estimated a significant total petrichor VOC emission (∼20 kg km^−2^ day^−1^), which is of the same order of magnitude as Israel’s reported anthropogenic VOC emissions (∼14.4 kg km^−2^ day^−1^) ^42^. The ozone formation potential (55 kg km^−2^ day^−1^) and SOA formation potential (1.36 kg km^−2^ day^−1^) associated with petrichor emissions are comparable with (∼25% lower) than anthropogenic VOC emissions in a highly urbanized area such as Los Angeles^43^. Compared to Los Angeles’s urban forest biogenic VOCs, petrichor VOCs show lower ozone formation but higher SOA formation, due to the dominance of 11 monoterpenoids rather than isoprene. Even without accounting for ∼6% built-up land cover (∼8-9% including inland water bodies) in Israel, these comparison values indicate that petrichor emissions from Israel may contribute to local atmospheric chemistry. The ultimate impact on oxidants and aerosols will depend on ambient NOx levels. Regional anthropogenic NOx emissions could be sufficient for producing ozone and SOA from petrichor VOC emissions in at least some parts of Israel. Soil microbes could be a significant NOx emission source at other locations, especially since these emissions strongly increase by rewetting^44^ similar to petrichor VOC emissions. The concurrent emission pulse of VOC and NOx after rewetting in drylands could stimulate cloud condensation nuclei production^45^, resulting in a coupling between biogenic emissions, aerosol, and precipitation.

Although the blend of VOC composing petrichor might vary with the climatic regions, the magnitude of emissions was similar across the aridity gradient. Given that drylands cover ∼half of Earth’s land surface^8,9^, and are expanding due to climate change, petrichor emissions may influence atmospheric oxidation capacity, aerosol formation and ozone chemistry at regional to global scale. Despite their short duration, these episodic but spatially extensive pulses could be relevant for atmospheric processes. Our results highlight rewetted dryland soils as a previously overlooked source of atmospheric reactivity, especially as climate change is expanding drylands globally.

### Conclusion

Petrichor emissions from desert soils, although episodic, but may cause strong biogeochemical events. The first rewetting pulse after a long period of drought, dominated by benzenoids and terpenoids, has the potential to impact atmospheric chemistry of SOA and tropospheric O3 formation. Terpenoids and sulfur VOCs are tied to microbial interactions and secondary metabolism. These compositional shifts may mirror desert microbial strategies or consequences of rapid resuscitation and have important implications for atmospheric oxidation capacity and aerosol formation. Under climate change, altered rainfall regimes will reshape not only the magnitude but also the chemistry of dryland VOC emissions, with implications for both ecosystem resilience and air quality.

## Materials and Methods

### Soil collection

Soil cores were collected at the end of the dry season (end of August) from a transect spanning ∼160 km in a north-south orientation across the Judea Hills and Negev Desert of Israel, established for sampling and representing an aridity gradient^13,14^ (Fig. 1). Soils were not hydrated for at least 3 months. Collection sites were located within four distinct climatic regions, each distinguished by annual precipitation: sub-humid (SH) shrubland (>300 mm yr^−1^), semi-arid (SA) grassland (∼200-300 mm yr^−1^), arid (A) desert (∼ 50-200 mm yr^−1^), and hyper-arid (HA) desert (<50 mm yr^−1^). Full data on soil moisture, temperature regimes, soil taxonomy order, soil properties and aridity index can be found in supplementary material (Fig. S1-2, Data S1, Table 1). Five replicate soil cores were obtained from two sampling sites in each climatic region (‘A’ and ‘B’, distant ∼2-10 km apart). Soil texture and soil water content (SWC) are provided in the supplements (Table S1).

To preserve soil structure and biocrust integrity, aluminum columns (56 mm × 40 mm; d × l) were inserted vertically into the soil until their upper edges were level with the biocrust surface. Cotton pads were placed on top of each column to protect the biocrust, and the column was carefully excavated and gently extracted from the soil. Then a second cotton pad was placed at the bottom, the pads were secured with aluminum foil, and the column was sealed with PET foils. The secured columns were transported to Helmholtz Munich (Germany) for analysis. Altogether 40 columns were analyzed (5 replicates × 2 sites × 4 climatic regions). Samples were stored under controlled conditions (21 ± 1 °C; 50 ± 10% RH) until analysis.

### Experimental design

Soil columns were rewetted in two simulated rain events. Rewetting was achieved by slowly dropping 25 mL of Milli-Q water in five sequential injections of 5 mL each at a rate of 1.5 mL min⁻¹ through an injection port and using a glass syringe. This corresponds to a rainfall of ∼10 mm, which is a realistic single precipitation event in Israel, where annual rainfall ranges from >600 mm in Mediterranean regions to <50 mm in the hyper-arid Negev Desert. The first rewetting followed >3 months of drought, and the second after 31 days of desiccation (Fig. S3). Three replicate soils were continuously monitored for 4 days after each rewetting event. Soils were incubated in 12 h light/dark cycles (135 ± 15 µmol m^−2^ s^−1^ PPFD) with temperatures of 28.4 ± 0.2°C (day) and 22.8 ± 0.1°C (night) and relative humidity of 11 ± 1% / 65 ± 2%, respectively (Fig. S4).

VOC fluxes were measured before and after the rewetting events under controlled environmental conditions using dynamic flow-through cuvettes^46,47^. The original glass cuvettes were replaced by ten 1 L odorless PET chambers to enable simultaneous measurements during simulated rainfall. Purified air (200 mL min^−1^) continuously flushed each cuvette, maintaining VOC mixing ratios constant and below 10 ppbv, and CO2 at 420 ppmv (Fig. S5). Outlet air was split to a proton-transfer-reaction quadrupole-mass-spectrometry (PTR-QMS) for online VOC analysis, a thermal desorption gas chromatography mass spectrometry (TD-GC-MS) for offline analysis, and an infrared gas analyser for CO2 and H2O. Cuvettes were switched automatically every 6 min, yielding a 1 h time resolution. The first 4 min of each cycle were discarded to avoid carry-over. The cuvettes were measured in randomized order. Full experimental details are provided in Method S2 and Figs S3-S6.

### VOC analysis (PTR-QMS)

Protonated VOCs were measured by PTR-QMS following established methods^48,49^ (detailed description in Method S3). A list of the monitored protonated masses and their chemical assignments is provided in Table S2. Geosmin was monitored at m/z 95 (fragment), and monoterpenes at m/z 137 (parent ion). The signal at m/z 71, typically associated with methyl vinyl ketone (MVK) and methacrolein (MACR), was not confirmed by GC-MS in our samples. Therefore, we tentatively attribute this ion to 1-octen-3-ol, which was identified by GC-MS. The VOCs collected on the GC-MS adsorbent tubes were analyzed using both untargeted and targeted approaches. Untargeted analysis was performed using MS-DIAL software^50^. Targeted identification of known volatiles was carried out using authentic reference standards^46,48,49,51–56^. The VOC fluxes were calculated as previously described^57^, based on ground area (m^2^).

### Gas exchange (CO2 and H2O) fluxes

Gas-exchange of CO2 and H2O were measured by IRGA (LI-850 LI-COR). Calibration of the instrument was achieved by certified standard gases (Air Liquide Deutschland GmbH, Düsseldorf, Germany) in the range of 0-2000 ppmv. CO2 and H2O fluxes were calculated as before^57^, using the soil surface area as the basis of the normalization. Positive fluxes of CO2 and H2O mean emissions from the soil to the atmosphere. The values were calculated individually for each mesocosm type and climate condition.

### Analysis of bacterial community

To characterize bacterial communities across the aridity gradient, we re-analysed a previously generated 16S rRNA amplicon dataset from the same climatic regions and sites^13^. Soil samples were collected from the same locations and separated into biocrust (0-2 cm) and topsoil (2-10 cm). DNA was extracted and 16S rRNA gene amplicon sequencing was performed as previously described^13^. We re-processed the sequence reads to generate updated community composition profiles for the present study.

### Assessment of potential impact on SOA and O3

The petrichor mol emission rates (ER) (pmol m^−2^ s^−1^) were converted to g m^−2^ h^−1^ using MW for each compound. Estimates of the maximum incremental reactivity (MIR) and SOA yields (Y_SOA) for each petrichor compound^43^ were used to estimate OFP and SOAP of each compound as:

OFP = ER x MIR

SOAP = ER x Y_SOA

### Statistical analysis

The VOC data was analyzed by multivariate data analysis, t-test and ANOVA. The principal component analysis (PCA) and orthogonal partial least square regression discriminant analysis (OPLS-DA) were performed using SIMCA-P v.13.0.3.0 (Umetrics, Umeå, Sweden) as previously described^46,52,53^. For multivariate analysis, data were preprocessed by log10 transformed, mean-centered, and Pareto scaled to conform to a normal distribution. Significant components were tested with the cross-validation method^58^ using a 99% confidence level on parameters and seven cross-validation groups. The significance of OPLS-DA was assessed using cross-validated ANOVA^58^ and was considered statistically significant at p < 0.05. Hierarchical clustering based on Spearman correlations, Kruskal-Wallis test, PERMANOVA and multiple test corrections (Benjamini-Hochberg procedure) were performed in R (R Core Team D, 2021). Differences in bacterial community composition across habitat types and climatic regions were assessed using permutation-based multivariate analyses on Hellinger-transformed 16S rRNA amplicon data, with tests for both centroid shifts and dispersion effects (see Methods S4 for details).

## Data availability

All data that support the findings of this study are publicly available on OSF (https://osf.io/jc65s) and on Sequence Read Archive under BioProject accession number PRJNA657906.

## Acknowledgments

We thank Elena Consorti for helping during the collection of VOC samples.

## Declaration of authorship

AG, OG, and HS conceived the study. BP and OG performed microbial analyses. MJ constructed oil maps. BP and HS performed field collections and soil analyses. AG and BW performed the experiments and analyzed the VOC/CO2 data. ABG studied the atmospheric impact. AG wrote the irst draft of the manuscript with the help of OG, ABG, JPS and MB. All authors contributed to the interpretation of the findings and edited and approved the manuscript.

