## Supplementary information for "Dryland soil rewetting induces strong VOC emissions with potential to form ozone and aerosols"

This file contains additional results and details on the study site, methods, data sources used in this study.

Table of Contents:

**Supplementary Figures**

Figure S1 - Soil characteristics

Figure S2 - Soil physicochemical properties

Figure S3 - Experimental timeline

Figure S4 - Air temperature

Figure S5 - Schematic of the dynamic flow-through cuvette enclosure system

Figure S6 - Soil cores

Figure S7 - Volatiles strongly correlating to rewetting

Figure S8 - Geosmin

Figure S9 - Correlation coefficients of VOCs to climatic regions

Figure S10 - VOC emission rates of the volatiles correlated with the climate regions

Figure S11 - Emission dynamics of oxygenated VOCs

Figure S12 - Emission dynamics of terpene-related VOCs

Figure S13 - Emission dynamics of sulfur-containing and other VOCs

Figure S14 - Correlation heatmap of PTR-QMS ion signals

Figure S15 - Correlation heatmap of VOC emission rates and $CO_2$ and $H_2O$ fluxes

Figure S16 - Petrichor-driven VOC mass, $O_3$ and SOA formation potential

**Supplementary Tables**

Table S1 - Soil texture and water content analyses

Table S2 - List of m/z measured with PTR-QMS

Table S3 - Microbial compositions (*Excel file*)

Table S4 - Soil bacterial community differences

Table S5 - Homogeneity of dispersion

Table S6 - Climatic region effect on bacteria composition

Table S7 - Soil type effect on bacteria composition

Table S8 - ID of VOCs (*Excel file*)

Table S9 - VOC emission rates (*Excel file*)

**Supplementary Methods**

Method S1 - Soil analysis

Method S2 - Experimental design

Method S3 - VOC analysis

Method S4 - Statistical analysis of bacterial diversity and community composition

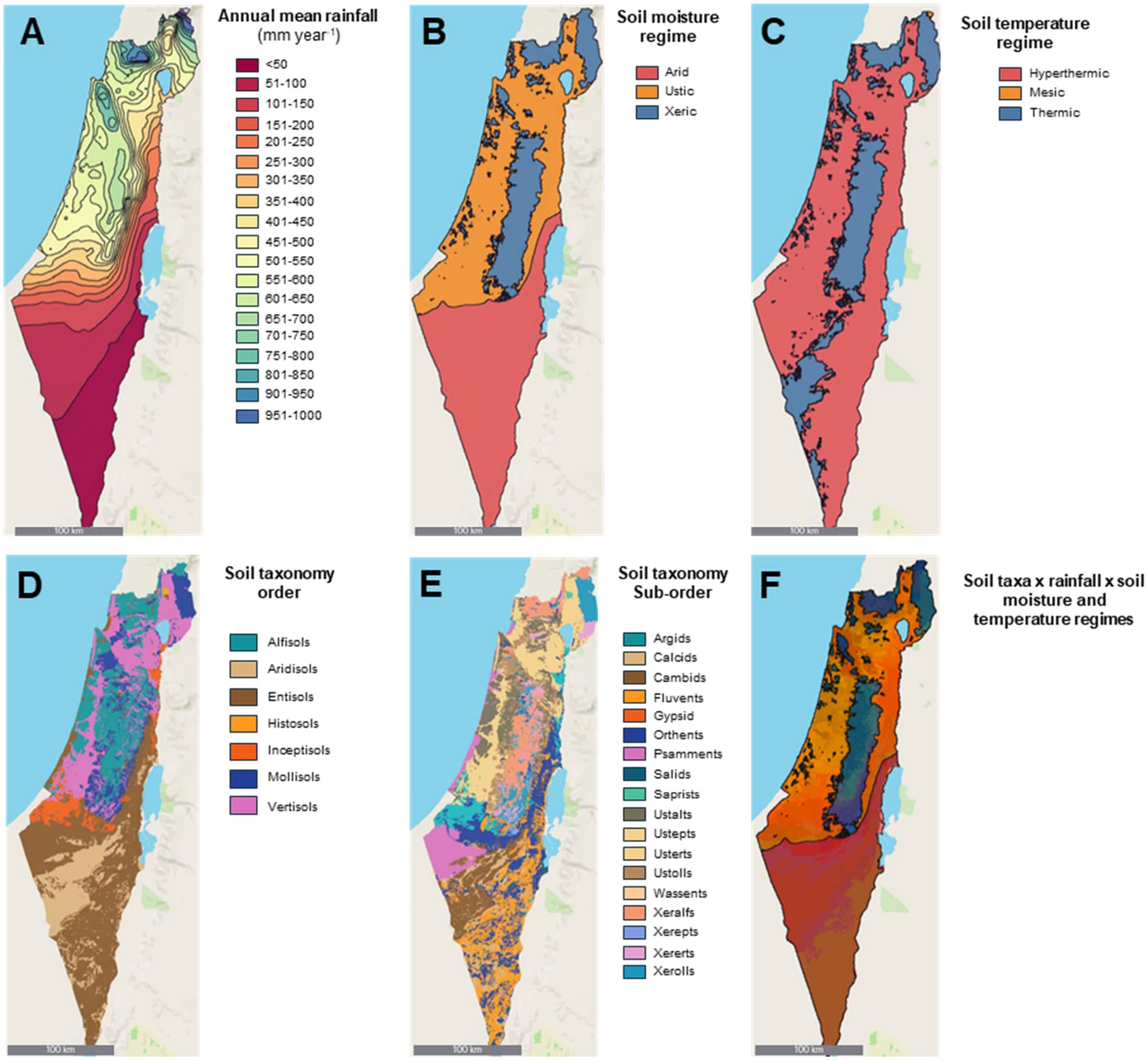


**Figure S1 - Soil characteristics of Israel along an aridity gradient.**

(**A**) Annual mean rainfall; (**B**) soil moisture regime; (**C**) soil temperature regime; (**D**) soil taxonomy order; (**E**) soil taxonomy suborder; (**F**) composite map overlaying panels A-D. Climate data and soil taxonomy are based on the USDA-NRCS Java Newhall Simulation Model (jNSM) and the USDA soil taxonomy system[1-3]. High-resolution datasets were merged to produce the maps in kepler.gl. Input data and GeoJSON (Geographic JavaScript Object Notation) files are available as Suppl. Data 1. Data retrieved from Israel ministry of agriculture (taxonomy, soil moisture and temperature data; https://data1-moag.opendata.arcgis.com) and Israel Meteorological service (annual mean rainfall data: https://ims.gov.il/en/ClimateAtlas).

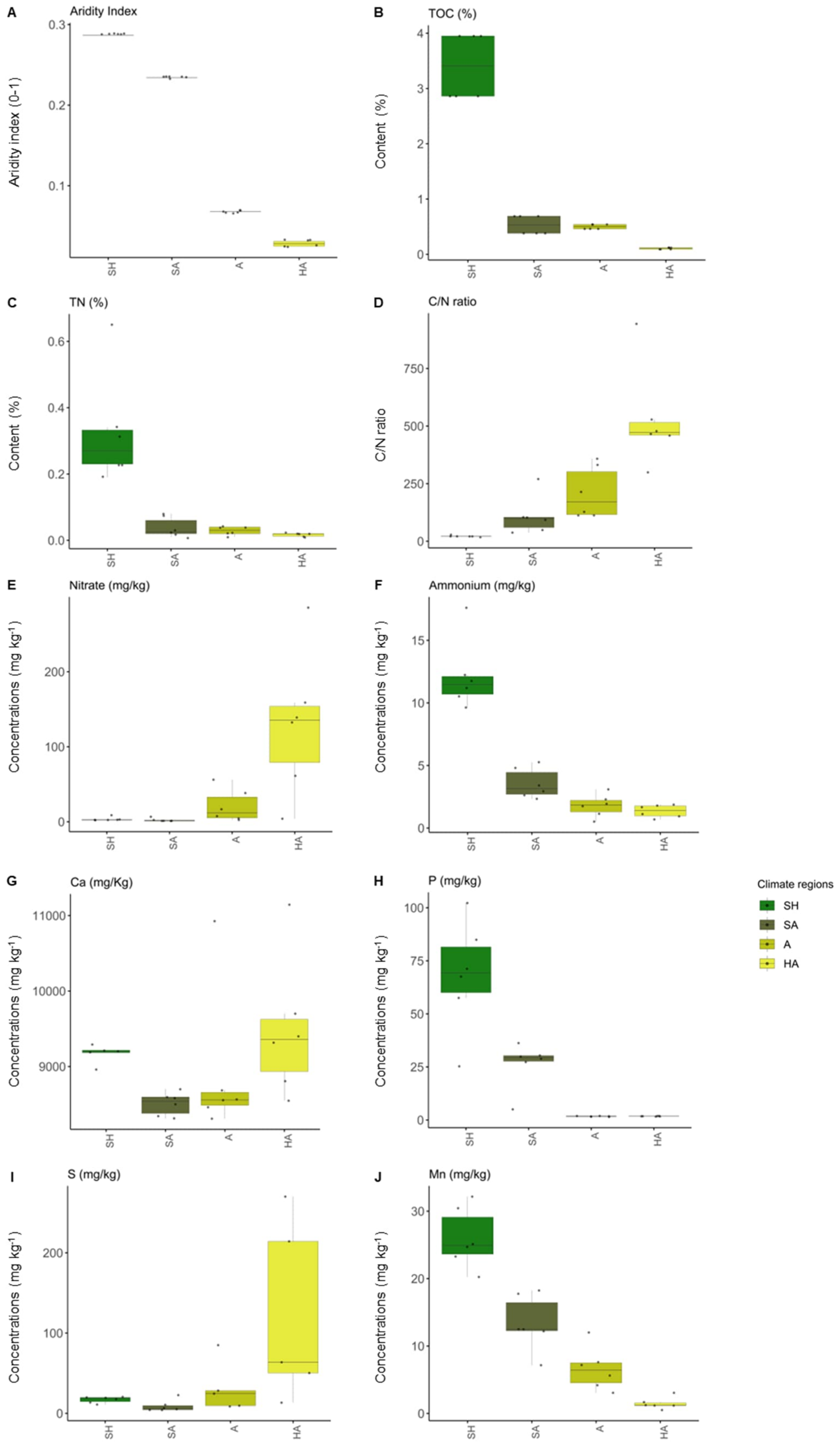


**Figure S2 - Soil physicochemical properties along the aridity gradient.**

(**A-J**) Boxplots of the aridity index, total organic carbon (TOC), total nitrogen (TN), carbon-to-nitrogen ratio (C/N), nitrate, ammonium, calcium (Ca), phosphorus (P), sulfur (S), and manganese (Mn) across the four climatic regions across the Judea Hills and Negev Desert of Israel: sub-humid (SH), semi-arid (SA), arid (A), and hyper-arid (HA). Data was extracted from [4], licensed under CC BY 4.0. A comprehensive analysis can be found in [4] .

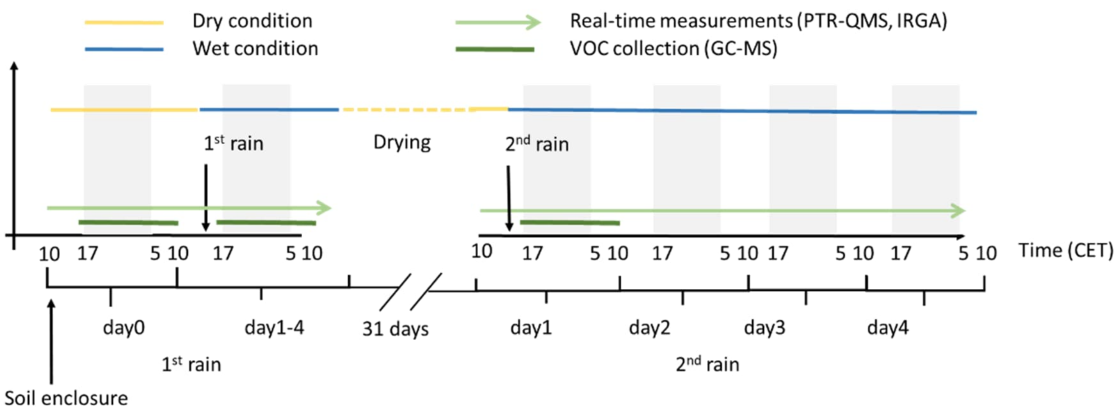


**Figure S3 - Experimental timeline for soil enclosure sampling, rain simulation, and VOC measurements.**

Schematic representation of the sampling schedule and simulated rainfall events during the experiment. Petrichor emissions were analyzed in cuvette-enclosed soils after the summer drought season (> 3 months), using TD-GC-MS and PTR-QMS following two simulated rain events, separated by a 31-day drying period. The study comprised an initial enclosure period (10-11 CET, day 0) and measurements under dry (control) conditions (yellow line), followed by the first rewetting event (16:30 CET; black arrow) and a 4-day monitoring period under wet conditions (blue line). VOC collection for GC-MS analysis began at 17:00 CET, 30 minutes after completion of the rain simulation, and continued for 17 hrs. After a drying phase of 31 days, a second rewetting was applied, followed by another 4-day wet period. Gray-shaded areas indicate the dark phase (12 hr, from 17 to 5, CET); the sampling time points for VOC collection using sorbent tubes and subsequent TD-GC-MS analysis are indicated by dark green lines. Real-time VOC measurements by PTR-QMS and $CO_2$/$H_2O$ monitoring by infrared gas analyzer (IRGA) were performed continuously throughout each observation period (light green arrows). Control measurements of petrichor emissions (i.e., under dry conditions) were collected the day prior to the rain simulation.

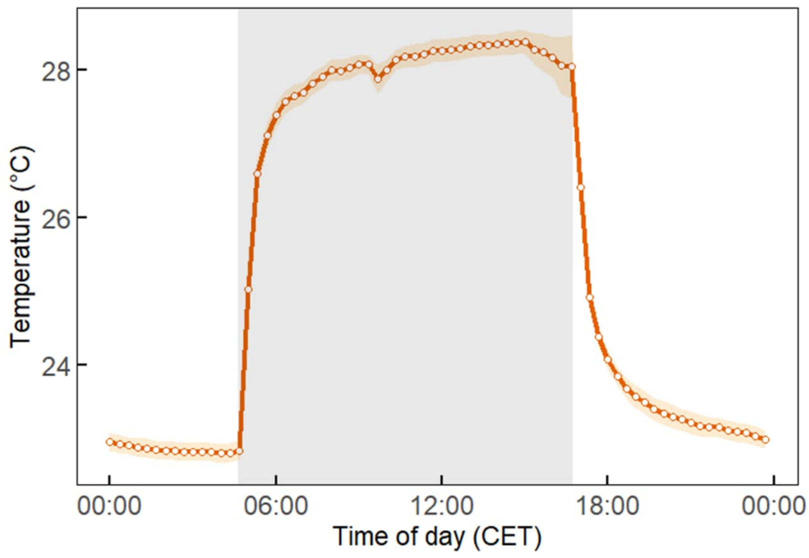


**Figure S4 - Air temperature inside the flow-through cuvettes during the VOC measurements.**

Mean (± 90% confidence interval) of 20-minute-averaged air temperature. The light was switched on between 5:00 and 17:00 CET (in gray).

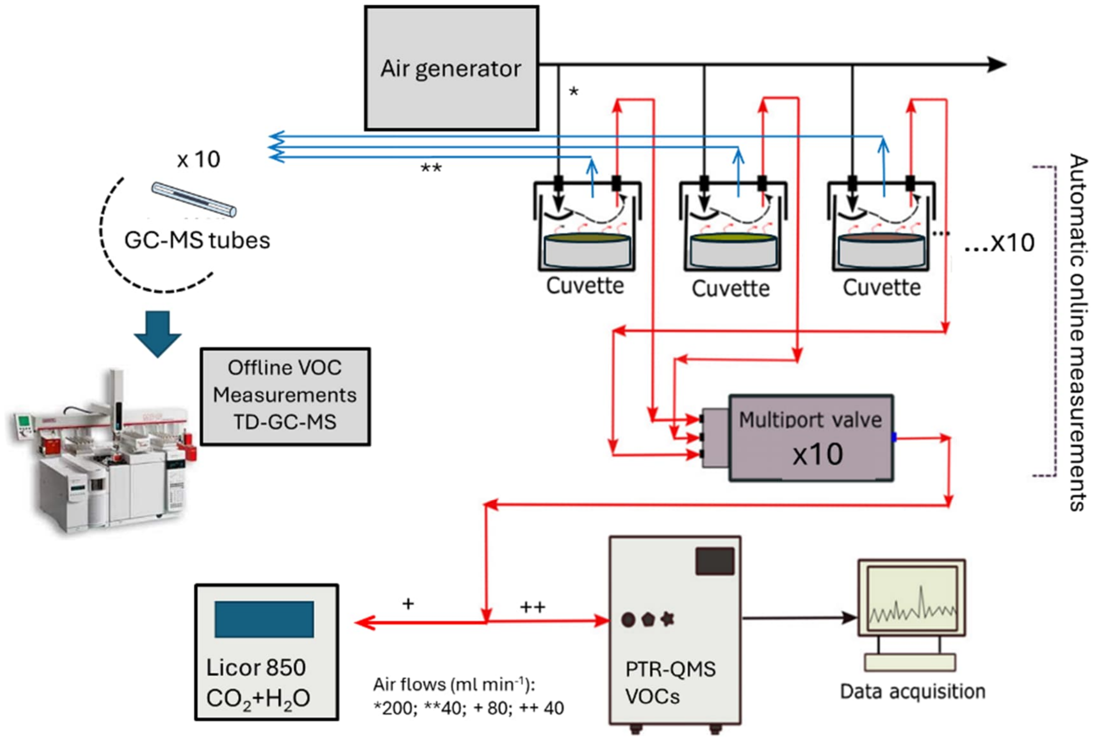


**Figure S5 - Schematic of the dynamic flow-through cuvette enclosure system for VOC measurements.**

The setup included ten cuvettes enclosing mesocosms that were continuously flushed with purified air (VOC < 10 ppb). Air exiting each cuvette was directed either to automated online analysis or to offline sampling. For online measurements, cuvette outflows were sequentially directed via an automated multiport valve to a proton transfer reaction quadrupole mass spectrometer (PTR-QMS) for real-time monitoring of VOCs and to a LI-COR 850 analyzer for $CO_2$ and $H_2O$ vapor measurements. Data was recorded continuously. For offline VOC analysis, air was collected onto sorbent cartridges and subsequently analyzed by thermal desorption-gas chromatography-mass spectrometry (TD-GC-MS).

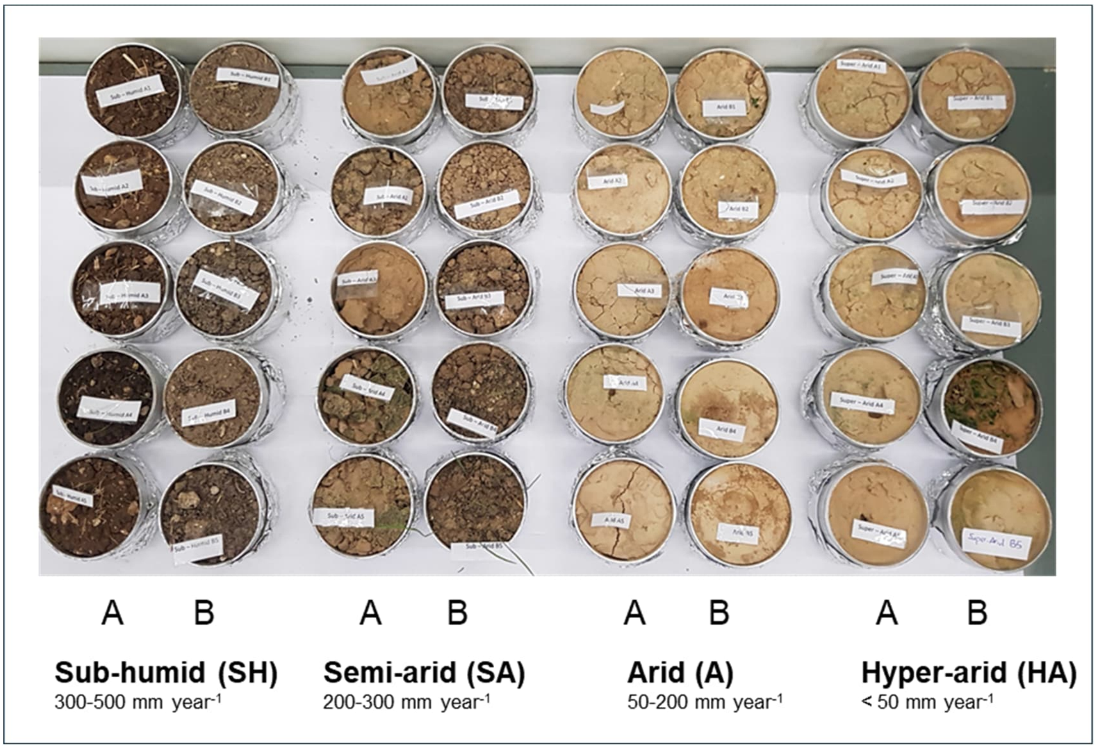


**Figure S6 - Soil cores at the end of the experiments.**

For each of the four climate regions (SH, sub-humid; SA, semi-arid; A, arid; HA, hyper-arid), two sites located 2-10 km apart (labeled ‘A’ and ‘B’) were selected, and five replicate cores were collected per site, yielding a total of 40 cores. This sampling design accounted for within-region heterogeneity and provided replicate soil samples for VOC analyses.

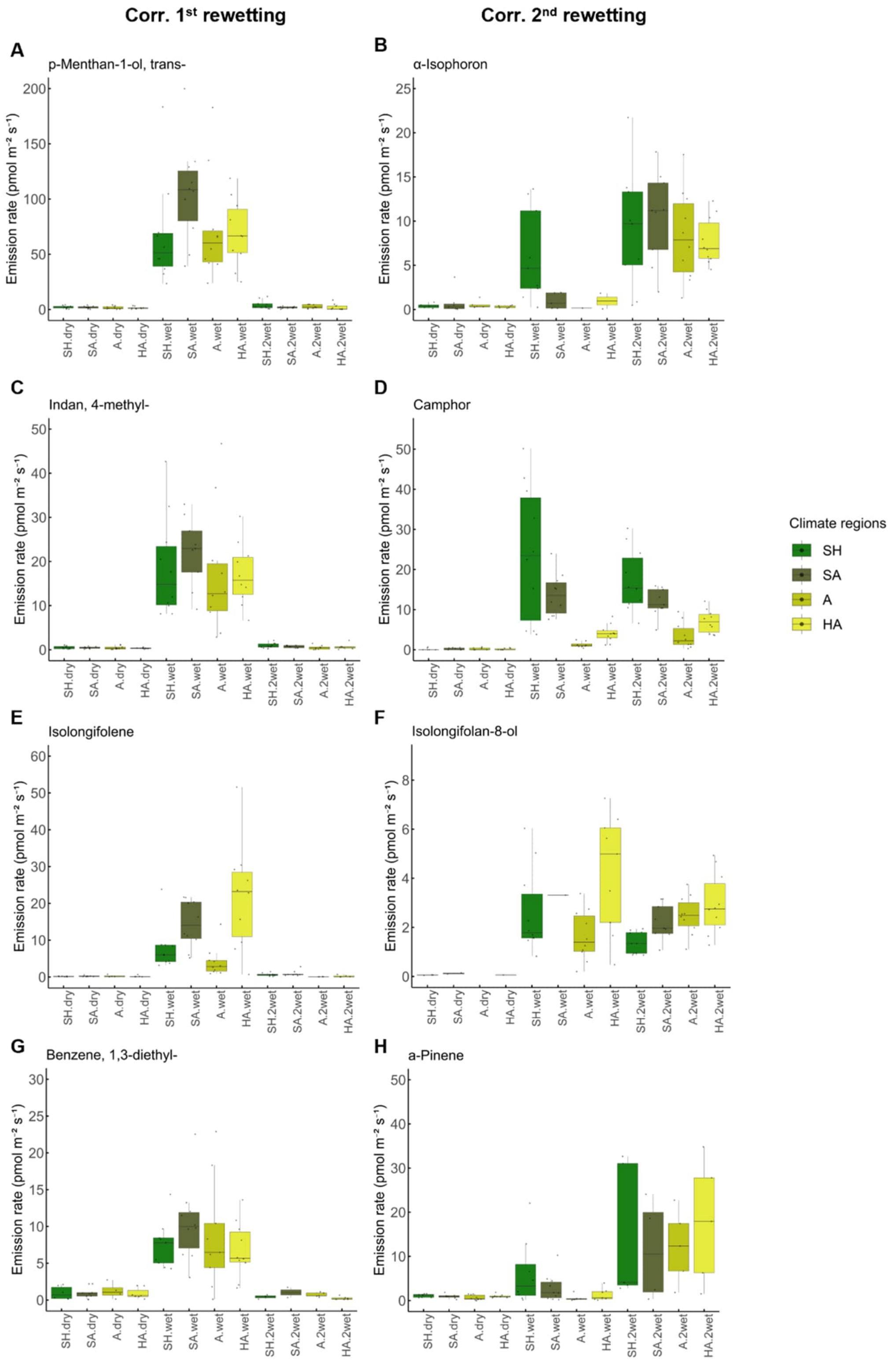


**Figure S7 - VOC emissions rates of the volatiles strongly correlated with the first (left) and second (right) rewetting episodes.**

Emission rates (pmol $m^{-2}$ $s^{-1}$) of (**A**) trans-p-menthan-1-ol, (**B**) α-isophorone, (**C**) 4-methyl-indan, (**D**) camphor, (**E**) isolongifolene, (**F**) isolongifolan-8-ol, (**G**) 1,3-diethyl-benzene, and (**H**) α-pinene, that showed strong positive correlations (OPLS, Fig. 2) with the first (left panels) and second (right panels) soil rewetting events across the aridity gradient. Box plots indicate median, interquartile range, and individual data points (n = 10 per climate region). Colours denote the four climatic regions: sub-humid (SH, dark green), semi-arid (SA, olive), arid (A, brown), and hyper-arid (HA, yellow). Treatments are labelled as ‘dry’ (before rewetting), ‘wet’ (first rewetting) and ‘2wet’ (second rewetting).

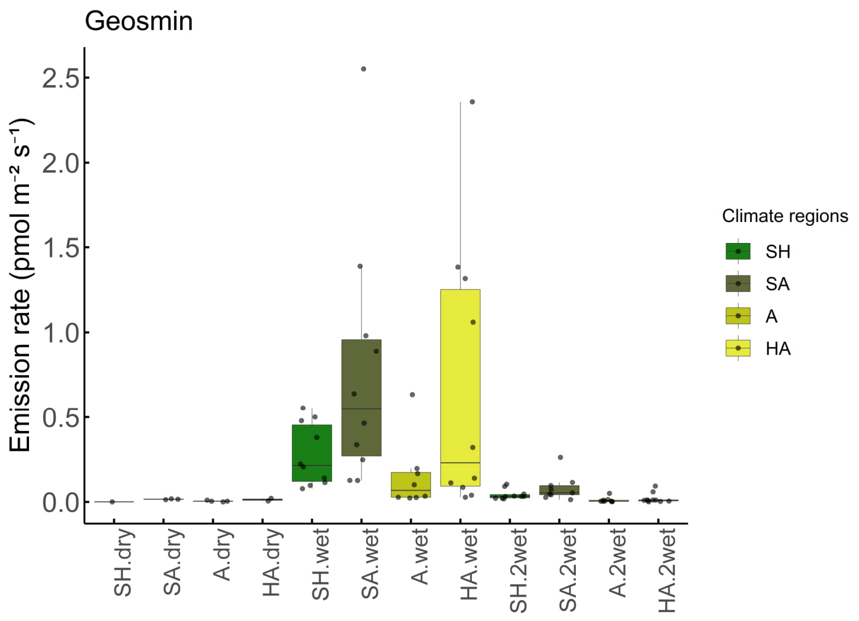


**Figure S8 - Geosmin emission rates.**

Emission rates (pmol $m^{-2}$ $s^{-1}$) were measured across soils from the four climatic regions along the aridity gradient under dry condition, first rewetting ('wet') occurred after a long period of field drought (> 3 months), and second rewetting ('2wet') occurred after a shorter desiccation period (31 days) in the laboratory. Box plots display medians, interquartile ranges, and individual data points (n = 10 per treatment). Colours indicate climatic regions: sub-humid (SH, dark green), semi-arid (SA, olive), arid (A, brown), and hyper-arid (HA, yellow).

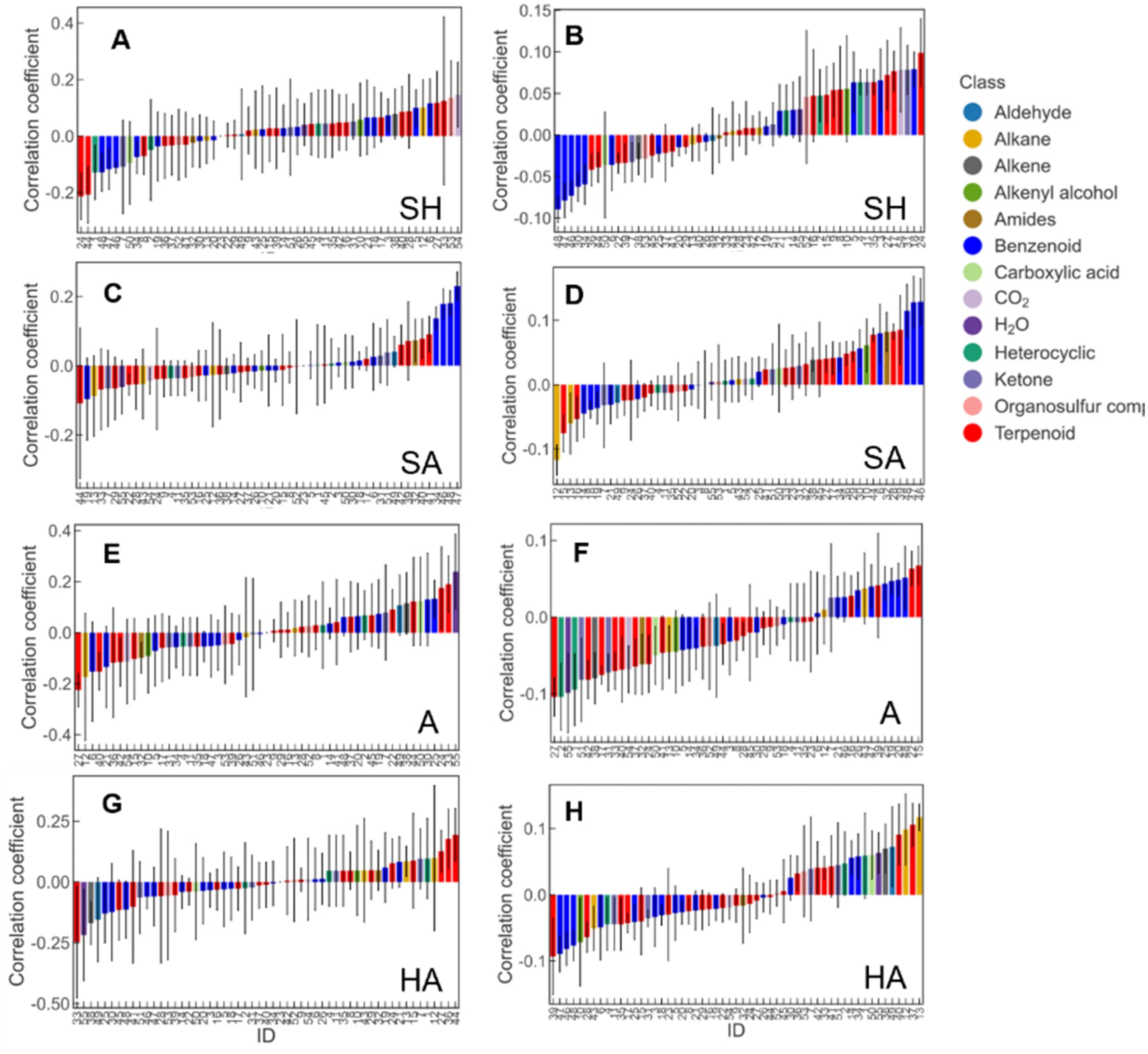


**Figure S9 - Correlation coefficients of individual VOCs derived from orthogonal partial least squares (OPLS) models corresponding to each of the four climatic regions along the aridity gradient.**

The correlation coefficients explain the group separation for: (**A-B**) sub-humid (SH), (**C-D**) semi-arid (SA), (**E-F**) arid (A), and (**G-H**) hyper-arid (HA) climate regions, under the two rewetting episodes. Bars show the mean correlation coefficients (± SD) of each VOC (compound ID, listed in Table S8) with the first (left panels) and second (right panels) soil rewetting events across the aridity gradient. Colours correspond to chemical classes as indicated in the legend. Positive coefficients indicate higher emissions from soils of a given climate region and compared to the others, whereas negative coefficients indicate lower emissions. The correlation coefficients correspond to the OPLS analysis shown in Figure 3.

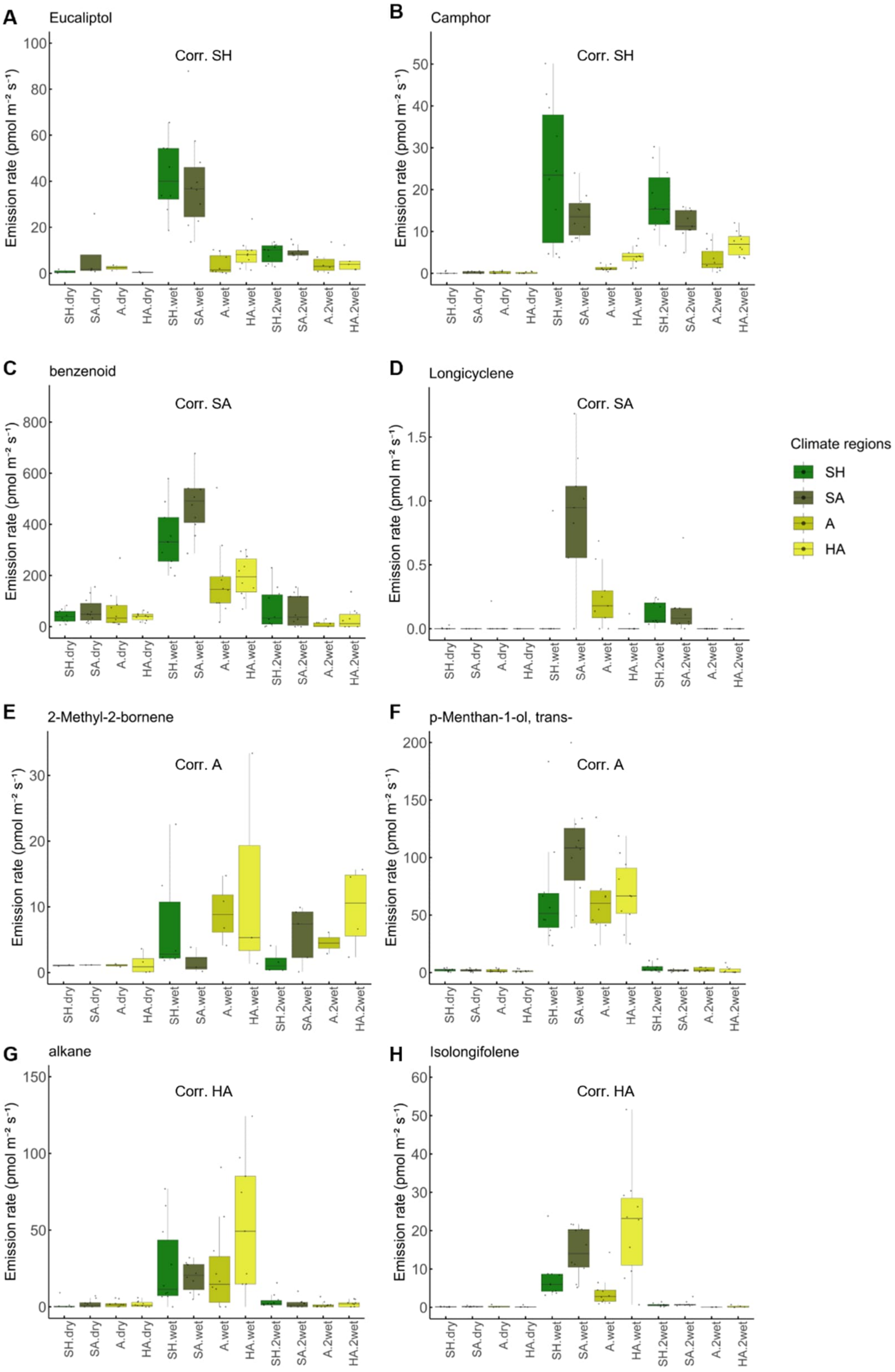


**Figure S10 - Emission rates of VOCs that correlated with the climate regions.**

Emission rates (pmol $m^{-2}$ $s^{-1}$) of VOCs that showed positive correlations (OPLS, Fig. 3) with soils from the four climatic regions along the aridity gradient. VOCs are grouped by the region where their emissions were most strongly correlated: SH region, (**A**) eucalyptol, (**B**) camphor; SA region, (**C**) total benzenoid emissions, (**D**) longicyclene; A region, (**E**) 2-methyl-2-bornene, (**F**) trans-p-menthan-1-ol; HA region, (**G**) total alkane emission, and (**H**) isolongifolene. Box plots indicate median, interquartile range, and individual replicates (n = 10 per climate region). Colours represent the four climate regions: sub-humid (SH, dark green), semi-arid (SA, olive), arid (A, brown), and hyper-arid (HA, yellow). Treatments are labelled as ‘dry’ (before rewetting), ‘wet’ (first rewetting) and ‘2 wet’ (second rewetting).

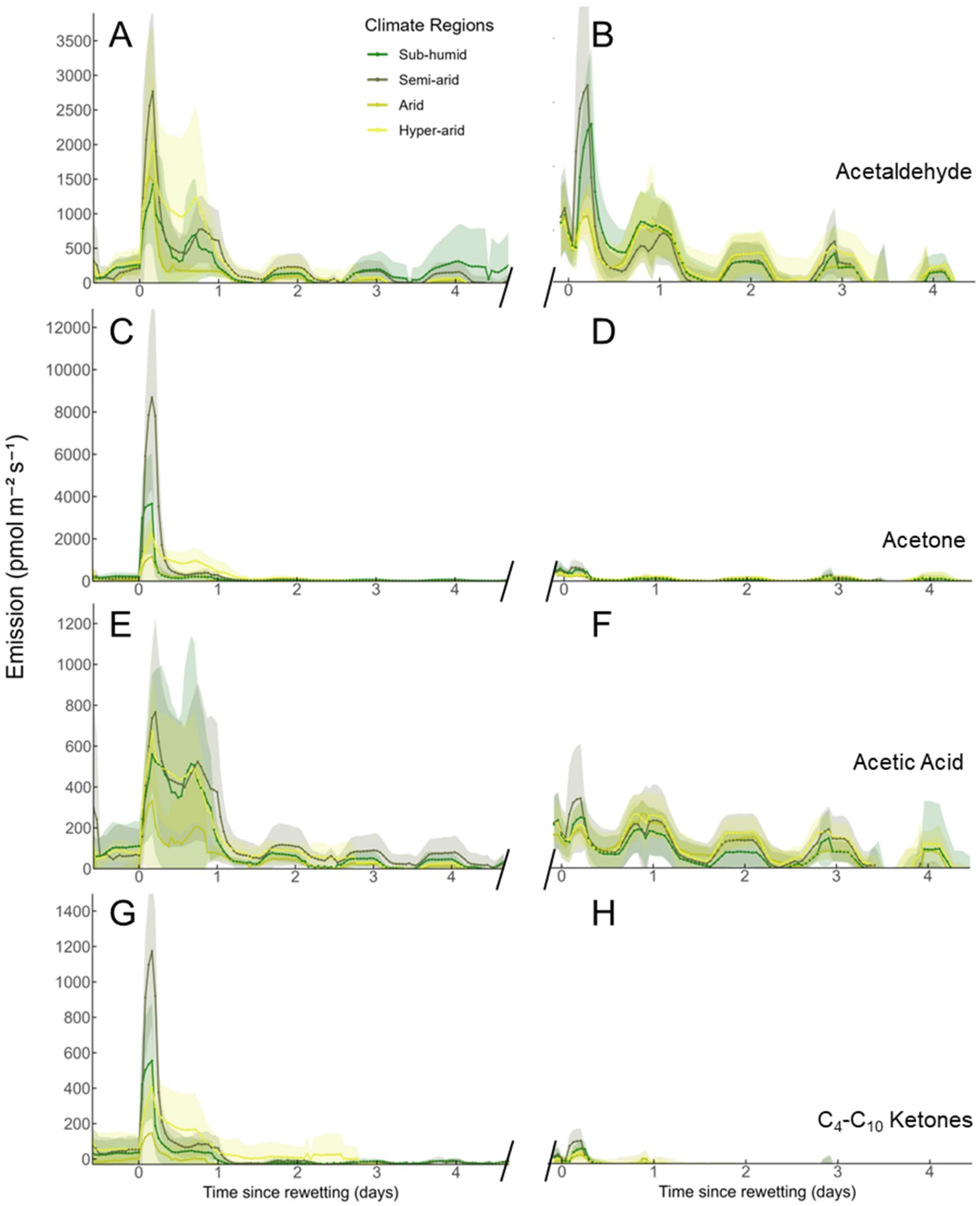


**Figure S11 - Emission dynamics of oxygenated VOCs in response to two rewetting events.**

Time series of VOC emission rates (normalized to ground area) measured by PTR-QMS are shown for acetaldehyde (**A-B**), acetone (**C-D**), acetic acid (**E-F**), and $C_4$–$C_{10}$ ketones (**G-H**). Panels on the left represent emissions following the first rewetting of dry soils, while panels on the right depict the response to a second rewetting event conducted after 31 days of desiccation. Soil cores originate from the four climate regions: sub-humid (SH); semi-arid (SA), arid (A), and hyper-arid (HA). Time is expressed relative to the onset of rewetting (day 0). Lines indicate mean fluxes and shaded areas represent 95% confidence intervals (n = 10).

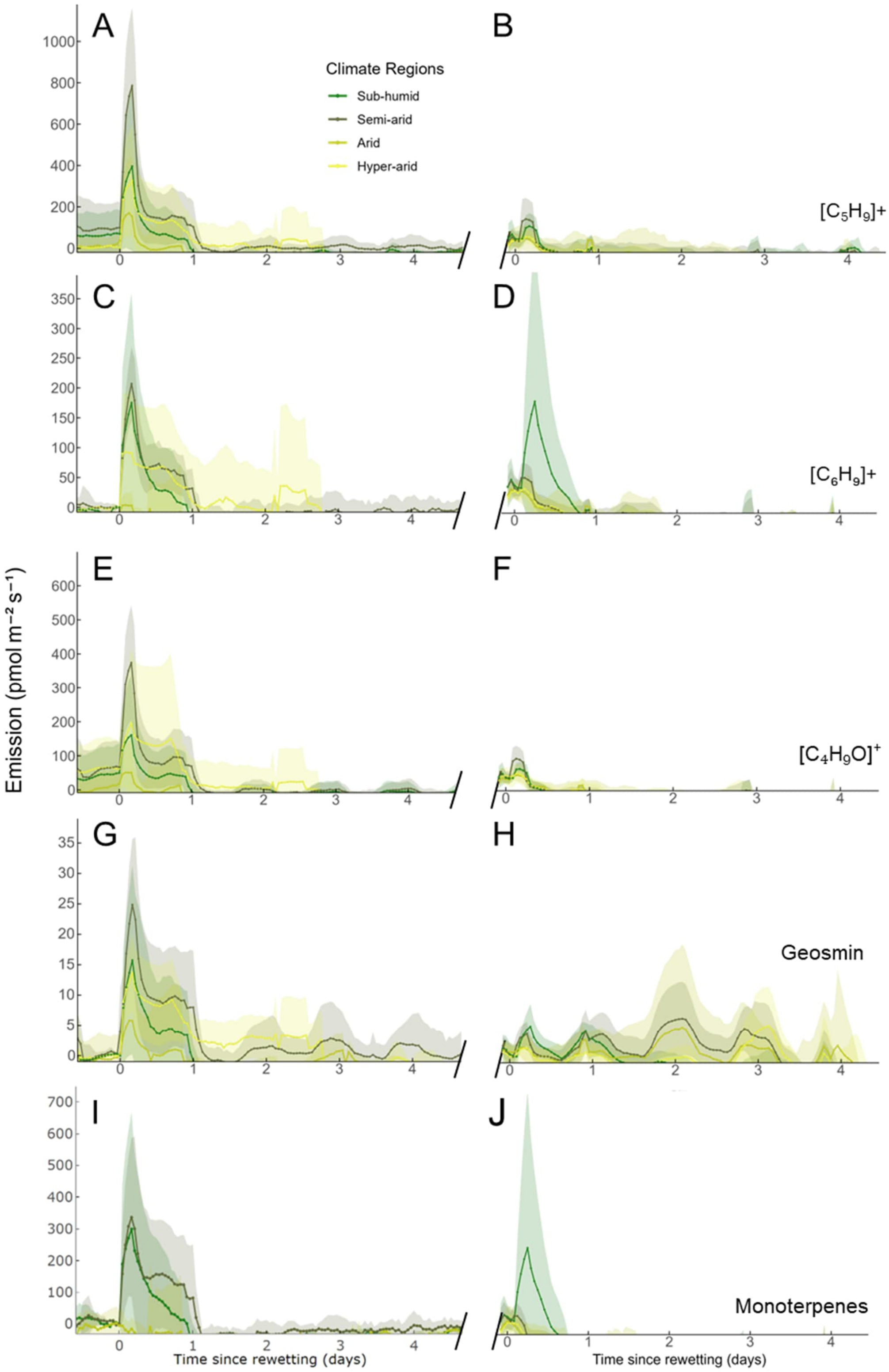


**Figure S12 - Emission dynamics of terpene-related VOCs in response to two rewettings.**

Time series of VOC emission rates (normalized to ground area) measured by PTR-QMS are shown for the monoterpenoid and sesquiterpenoid $C_5H_9^+$ (A-B) and $C_6H_9^+$ (C-D) fragments, $C_4H_9O^+$ (E-F), geosmin (G-H) and monoterpenes (I-J). Panels on the left represent emissions following the first wetting of dry soils, while panels on the right depict the response to a second wetting event conducted after 31 days of desiccation. Soil cores (~4cm deep) originate from the four climate regions sub-humid (SH), semi-arid (SA), arid (A), and hyper-arid (HA). Time is expressed relative to the onset of rewetting (day 0). Lines indicate mean fluxes and shaded areas represent 95% confidence intervals (n = 10). Putative identifications of the molecular formula are given in Supplementary Table S2.

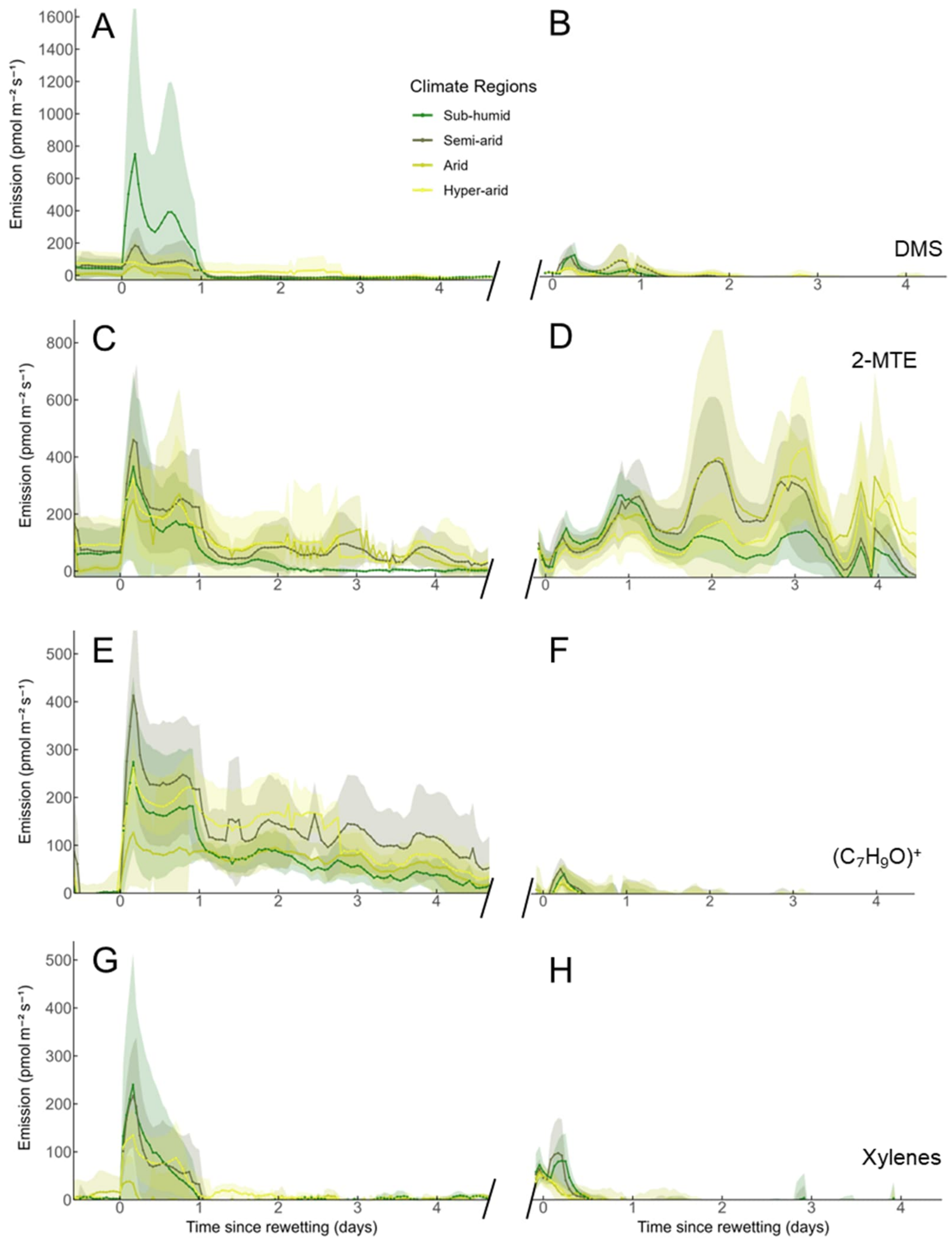


**Figure S13 - Emission dynamics of sulfur-containing and other VOCs in response to two rewetting events.**

Time series of VOC emission rates (normalized to ground area) measured by PTR-QMS are shown for dimethyl sulfide (DMS) (**A-B**), 2-methylthioethanol (2-MTE) (**C-D**), benzenoid and aromatic alcohols at $C_7H_9O^+$ (**E-F**), and xylenes (**G-H**). Panels on the left represent emissions following the first rewetting of dry soils, while panels on the right depict the response to a second rewetting event conducted after 31 days of desiccation. Soil cores originate from the four climate regions: sub-humid (SH), semi-arid (SA), arid (A), and hyper-arid (HA). Time is expressed relative to the onset of rewetting (day 0). Lines indicate mean fluxes, and shaded areas represent 95% confidence intervals (n = 10). Putative identifications of the molecular formula are given in Supplementary Table S2.

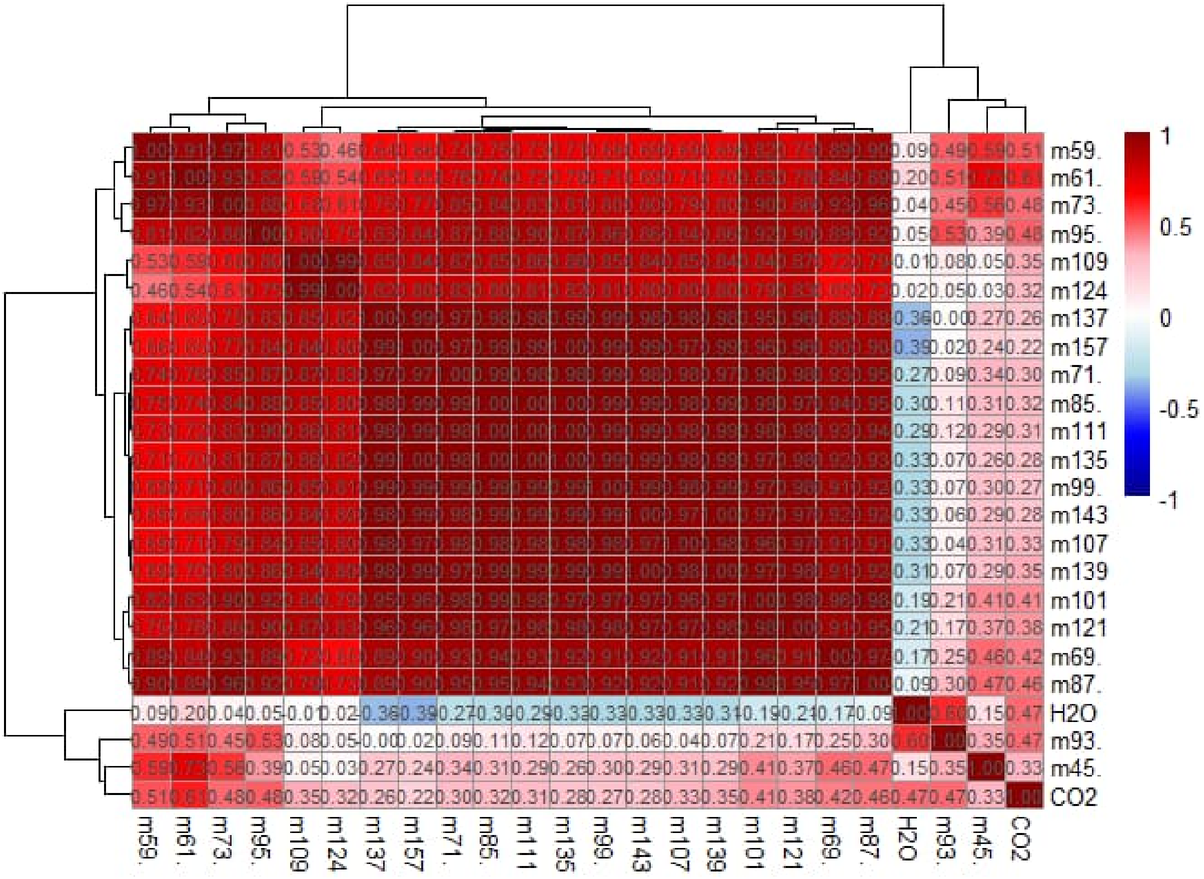


**Figure S14 - Clustered Spearman correlation heatmap of PTR-QMS ion signals and data on $CO_2$, and $H_2O$ fluxes.**

Spearman's rank correlation coefficients (ρ) among VOC fluxes measured by PTR-MS and $CO_2$ and $H_2O$ fluxes measured by infrared gas analysis (IRGA). Hierarchical clustering was based on spearman correlation. The color scale indicates strong positive (red) to strong negative (blue) correlations, with color intensity proportional to |ρ|. Dendrograms indicate similarity relationships among ion signal fluxes. The identified ion masses ('m') are listed in Suppl. Table S8. Abbr. of compounds mentioned in the text: m45: acetaldehyde; m59: acetone; m61: acetic acid; m95: geosmin fraction; m107/m109: benzenoids; m137: monoterpenes

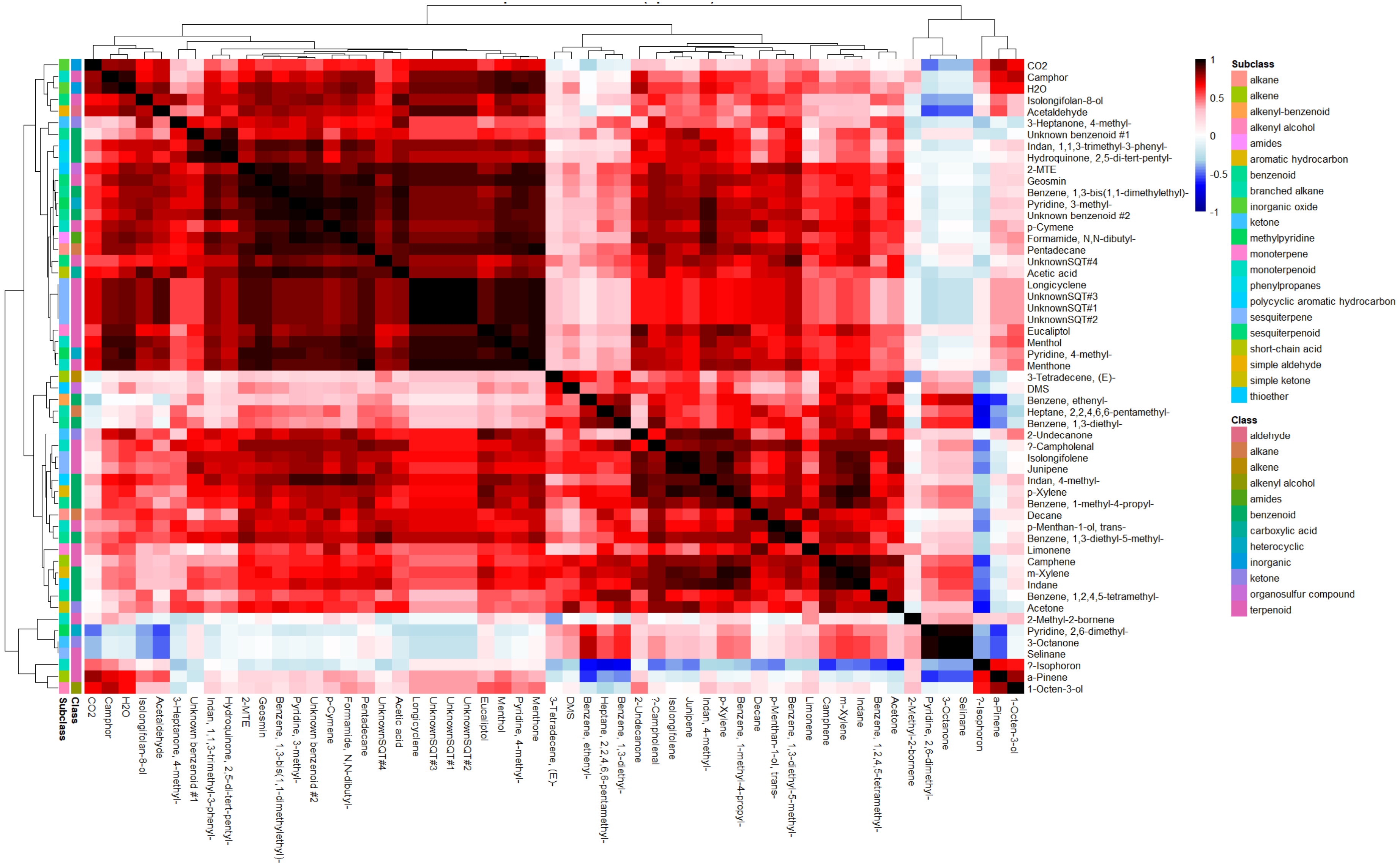


**Figure S15 - Clustered Spearman correlation heatmap of VOC emission rates and $CO_2$ and $H_2O$ fluxes across all the samples.**

Pairwise Spearman's rank correlation coefficients (ρ) between individual VOC emission rates measured by GC-MS and PTR-QMS and concurrent $CO_2$ and $H_2O$ fluxes measured by IRGA. Red colors indicate positive correlations and blue colors negative correlations. Dendrograms show the similarity structure among flux variables.

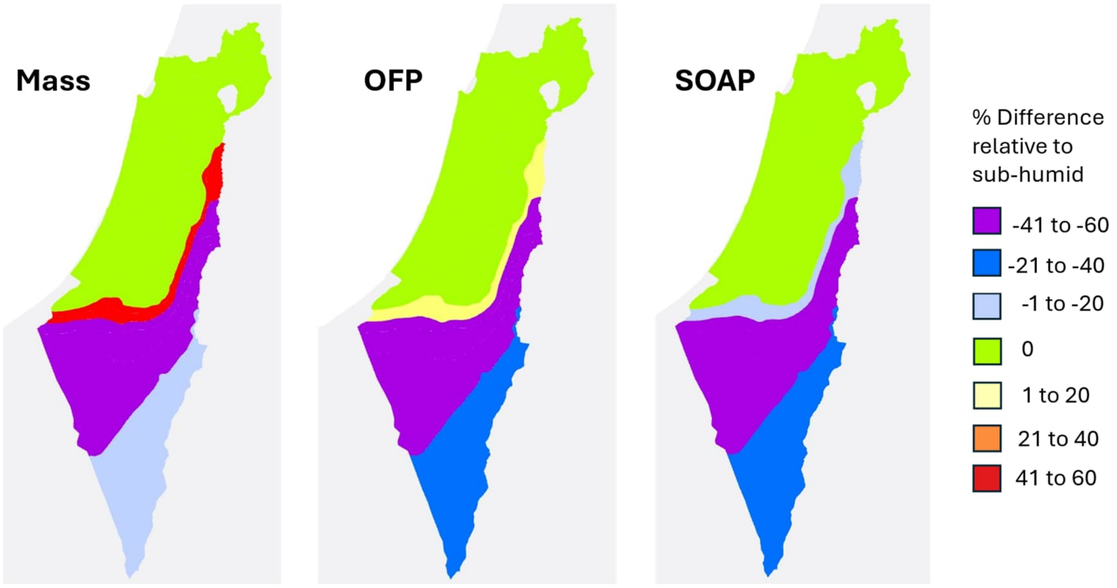


**Figure S16 - Petrichor-driven VOC mass, $O_3$ and SOA formation potential in Israel.**

Ozone-forming potential (OFP) and secondary organic aerosol potential (SOAP) were calculated from climate-region-specific petrichor VOC fluxes measured the day after rewetting, using maximum incremental reactivity (MIR) and SOAP weighting factors for each compound. The figure shows the relative mass OFP and SOAP formation to the sub-humid region related to post-rewetting petrichor emissions.

### Supplementary Tables

**Table S1 - Soil texture and water content across the aridity gradient.**

Soil texture and soil water content (SWC; % g $H_2O$ $g^{-1}$ dry soil) of samples collected along the aridity gradient, determined by sedimentation analysis (Method 1; n = 3 ± s.d.).

| | **Sand (%)** | **Silt (%)** | **Clay (%)** | **SWC (%)** |
|---|---|---|---|---|
| Sub-humid: | 50.0 ± 2.9 | 15.5 ± 2.0 | 34.5 ± 0.5 | 4.5 ± 0.85 |
| Semi-arid: | 56.3 ± 0.5 | 14.6 ± 1.4 | 29.2 ± 1.3 | 1.7 ± 0.33 |
| Arid: | 50.8 ± 2.3 | 13.1 ± 1.5 | 36.1 ± 1.6 | 1.3 ± 0.36 |
| Hyper-arid: | 44.4 ± 2.9 | 29.6 ± 1.6 | 25.9 ± 1.4 | 1.1 ± 0.11 |

**Table S2 - List of m/z values measured with proton-transfer-reaction quadrupole mass spectrometry (PTR-QMS).**

The ‘Compound’ column reports VOCs identified with high confidence based on supporting GC-MS data and known emission sources. The ‘Tentative identification’ column provides additional or alternative VOCs that may contribute to each m/z, assigned based on mass spectral interpretation and literature references. Abbreviations and protonated chemical formulas correspond to the assigned compound or compound class. Tentative assignments may include contributions from isobaric, co-eluting, or fragment ions not specified in the table; see main text for discussion of potential interferences and assignment confidence.

| m/z | Compound | Tentative identification | Abbreviation | Formula |
|---|---|---|---|---|
| 45 | Acetaldehyde | | Aceal | $C_2H_5O^+$ |
| 59 | Acetone | | Ace | $C_3H_7O^+$ |
| 61 | Acetic acid | | AA | $C_2H_4O_2^+$ |
| 63 | Dimethyl sulfide | | DMS | $C_2H_7S^+$ |
| 69 | $C_5H_9^+$ | Fragments of: alkenes / monoterpenoids / sesquiterpenoids | m/z 69 | $C_5H_9^+$ |
| 71 | $C_4H_9O^+$ | 1-octen-3-ol / methyl vinyl ketone (MVK) / methacrolein (MACR) | m/z 71 | $C_4H_9O^+$ |
| 81 | $C_6H_9^+$ | 1-octen-3-ol + fragment of monoterpenes, sesquiterpenes, geosmin, and various unsaturated $C_6$ alcohols or aldehydes | m/z 81 | $C_6H_9^+$ |
| 93 | 2-Methylthioethanol | | 2-MTE | $C_3H_9OS$ |
| 85+87 | $C_4$-$C_{10}$ Ketones | | | $C_4H_5O_2^+$ |
| 95+149+183 | Geosmin | | Geo | $C_{12}H_{23}O^+$ |
| 107 | Xylenes | | Xyl | $C_8H_{11}^+$ |
| 109 | $C_7H_9O^+$ | Benzenoid and aromatic alcohols | m/z 109 | $C_7H_9O^+$ |
| 137 | Monoterpenes | | MT | $C_{10}H_{17}^+$ |

**Table S3 - Bacterial community composition and 16S rRNA gene amplicon abundances for all samples.**

The table contains the full ASVs by sample matrix used for community analyses. Each row represents a single ASV, with taxonomic assignments. Each subsequent column corresponds to a sample. Cell values are the raw read counts per ASV per sample prior to any transformation (see .csv file).

**Table S4 - Soil bacterial community differences evaluated using PERMANOVA on the 16S rRNA gene amplicons.**

Statistical results of permutation-based multivariate analysis of variance (PERMANOVA) on Hellinger-transformed 16S rRNA gene amplicon profiles (Bacteria only). ASV counts were converted to relative abundances per sample, Hellinger-transformed, and analyzed using Euclidean distances in multivariate space. Shown for each factor are the F-statistic (*F*), proportion of variance explained ($R^2$), permutation-based *p*-value (999 permutations), and degrees of freedom (df) between and within soil type or climatic region.

| Factor | F | $R^2$ | p-value | df between | df within |
|---|---|---|---|---|---|
| Soil type (Biocrust, Topsoil) | 3.9376 | 0.07885 | 0.001 | 1 | 46 |
| Climatic region (Sub-Humid, Semi-Arid, Arid, Hyper-Arid) | 5.3731 | 0.26812 | 0.001 | 3 | 44 |

**Table S5 – Homogeneity of dispersion within groups (PERMDISP analogue)**

Distance-based statistical test results for homogeneity of multivariate dispersion applied to the Hellinger-transformed 16S rRNA gene amplicon data (Bacteria only). For each factor (soil type and climatic region), samples were projected into Hellinger space, group centroids were calculated, and the Euclidean distance of each sample to its group centroid was used as a measure of within-group dispersion. The table reports the F-statistic (F), permutation based p-value (999 permutations), degrees of freedom (df) between and within soil type or climatic region, and the mean distance to centroid for each group. These results indicate whether PERMANOVA effects reflect differences in group centroids (location) rather than being solely driven by unequal within-group variability (dispersion).

| **Factor** | **F** | ***p*-value** | **df between** | **df within** | **Mean dist Biocrust** | **Mean dist Topsoil** | **Mean dist Sub-Humid** | **Mean dist Semi-Arid** | **Mean dist Arid** | **Mean dist Hyper-Arid** |
|---|---|---|---|---|---|---|---|---|---|---|
| Soil type | $4.35E^{-07}$ | 1 | 1 | 46 | 0.7868 | 0.7868 | | | | |
| Climatic region | 11.37014 | 0.001 | 3 | 44 | | | 0.6002 | 0.7296 | 0.7030 | 0.7621 |

**Table S6 - Climatic region effect (Kruskal-Wallis test) on orders within the different soil types (biorust and topsoil).**

Results of the non-parametric tests for the effect of climatic region on the relative abundances of dominant bacterial orders within each soil type. ASV counts were aggregated to order level, converted to relative abundances per sample, and orders with a mean relative abundance ≥ 1% across all samples were retained for analysis. For each order and soil type, the table reports the Kruskal-Wallis H statistic, the associated p-value, and the Benjamini-Hochberg FDR-corrected *q*-value (based on 4-group comparisons across climatic regions). Orders with $q < 0.05$ are considered to exhibit significant shifts in relative abundance along the aridity gradient, corresponding to the patterns shown in the order-level relative abundance figure (Figure 1).

| Soil type | Order | H | p | p_adj |
|---|---|---|---|---|
| Biocrust | 0319-7L14 | 6.7481 | 0.0804 | 0.1048 |
| | Acetobacterales | 7.2797 | 0.0635 | 0.0896 |
| | Burkholderiales | 11.1732 | 0.0108 | 0.0203 |
| | Chitinophagales | 6.6225 | 0.085 | 0.1062 |
| | Chthoniobacterales | 11.9609 | 0.0075 | 0.0161 |
| | Cyanobacteriales | 13.6565 | 0.0034 | 0.0096 |
| | Cytophagales | 8.6304 | 0.0346 | 0.0547 |
| | Frankiales | 13.2312 | 0.0042 | 0.0096 |
| | Gaiellales | 10.6877 | 0.0135 | 0.0231 |
| | Gemmatimonadales | 2.1159 | 0.5487 | 0.5487 |
| | IMCC26256 | 7.3413 | 0.0618 | 0.0896 |
| | Isosphaerales | 18.1152 | 0.0004 | 0.0052 |
| | Kallotenuales | 14.8543 | 0.0019 | 0.0075 |
| | KD4-96 | 13.2399 | 0.0041 | 0.0096 |
| | Longimicrobiales | 13.358 | 0.0039 | 0.0096 |
| | Micrococcales | 5.6783 | 0.1284 | 0.1431 |
| | Micromonosporales | 15.2326 | 0.0016 | 0.0075 |
| | Microtrichales | 14.7036 | 0.0021 | 0.0075 |
| | Propionibacteriales | 3.4014 | 0.3338 | 0.3453 |
| | Pseudonocardiales | 17.6478 | 0.0005 | 0.0052 |
| | Pyrinomonadales | 11.3007 | 0.0102 | 0.0203 |
| | Rhizobiales | 17.7906 | 0.0005 | 0.0052 |
| | Rhodobacterales | 4.9377 | 0.1764 | 0.189 |
| | Rubrobacterales | 10.6348 | 0.0139 | 0.0231 |
| | Solirubrobacterales | 5.671 | 0.1288 | 0.1431 |
| | Sphingomonadales | 7.2022 | 0.0657 | 0.0896 |

| | | | | |
|---|---|---|---|---|
| | Tepidisphaerales | 14.5551 | 0.0022 | 0.0075 |
| | Thermomicrobiales | 6.1007 | 0.1068 | 0.1282 |
| | uncultured | 15.4014 | 0.0015 | 0.0075 |
| | Vicinamibacterales | 15.6804 | 0.0013 | 0.0075 |
| Topsoil | 0319-7L14 | 13.5468 | 0.0036 | 0.0135 |
| | Acetobacterales | 2.5182 | 0.472 | 0.5244 |
| | Burkholderiales | 12.3736 | 0.0062 | 0.0166 |
| | Chitinophagales | 3.3503 | 0.3407 | 0.4088 |
| | Chthoniobacterales | 11.76 | 0.0083 | 0.019 |
| | Cyanobacteriales | 6.3534 | 0.0956 | 0.1594 |
| | Cytophagales | 5.9877 | 0.1122 | 0.1772 |
| | Frankiales | 4.869 | 0.1816 | 0.2612 |
| | Gaiellales | 4.1398 | 0.2468 | 0.3219 |
| | Gemmatimonadales | 4.4782 | 0.2142 | 0.2921 |
| | IMCC26256 | 17.1415 | 0.0007 | 0.0066 |
| | Isosphaerales | 16.396 | 0.0009 | 0.0071 |
| | Kallotenuales | 20.6949 | 0.0001 | 0.0027 |
| | KD4-96 | 12.6391 | 0.0055 | 0.0165 |
| | Longimicrobiales | 12.2338 | 0.0066 | 0.0166 |
| | Micrococcales | 0.8136 | 0.8462 | 0.8462 |
| | Micromonosporales | 15.1459 | 0.0017 | 0.0073 |
| | Microtrichales | 15.1675 | 0.0017 | 0.0073 |
| | Propionibacteriales | 3.498 | 0.321 | 0.4013 |
| | Pseudonocardiales | 2.3213 | 0.5084 | 0.5448 |
| | Pyrinomonadales | 13.1815 | 0.0043 | 0.0142 |
| | Rhizobiales | 11.4453 | 0.0095 | 0.0205 |
| | Rhodobacterales | 4.8536 | 0.1828 | 0.2612 |
| | Rubrobacterales | 1.3262 | 0.7229 | 0.7479 |
| | Solirubrobacterales | 10.1029 | 0.0177 | 0.0332 |
| | Sphingomonadales | 2.9292 | 0.4027 | 0.4646 |
| | Tepidisphaerales | 9.2699 | 0.0259 | 0.0457 |
| | Thermomicrobiales | 10.6752 | 0.0136 | 0.0272 |
| | uncultured | 19.8796 | 0.0002 | 0.0027 |
| | Vicinamibacterales | 15.6892 | 0.0013 | 0.0073 |

**Table S7 - Soil type effect (Mann-Whitney U test) for each climatic region.**

Pairwise differences in the relative abundance of dominant bacterial orders between soil types within each climatic region. ASV counts were aggregated to the order level, converted to relative abundances per sample, and orders with mean relative abundance ≥ 1% across all samples were retained for analysis. For each order and climatic region, the table reports the Mann–Whitney U statistic, the associated p-value, and the Benjamini-Hochberg FDR-corrected *q*-value comparing biocrust vs topsoil. Orders with $q < 0.05$ are considered to differ significantly between soil types within that region, highlighting habitat-specific enrichment patterns superimposed on the aridity gradient.

| Climatic region | Order | U | p | p_adj |
|---|---|---|---|---|
| Arid | Solirubrobacterales | 0 | 0.0022 | 0.0093 |
| | Cyanobacteriales | 36 | 0.0022 | 0.0093 |
| | Gaiellales | 0 | 0.0022 | 0.0093 |
| | Acetobacterales | 36 | 0.0022 | 0.0093 |
| | Vicinamibacterales | 0 | 0.0022 | 0.0093 |
| | KD4-96 | 0 | 0.0022 | 0.0093 |
| | 0319-7L14 | 0 | 0.0022 | 0.0093 |
| | Longimicrobiales | 35 | 0.0043 | 0.0118 |
| | Pseudonocardiales | 1 | 0.0043 | 0.0118 |
| | Microtrichales | 1 | 0.0043 | 0.0118 |
| | Rhodobacterales | 35 | 0.0043 | 0.0118 |
| | Cytophagales | 34 | 0.0087 | 0.0216 |
| | IMCC26256 | 3 | 0.0152 | 0.0350 |
| Hyper-Arid | Gaiellales | 0 | 0.0022 | 0.0325 |
| | Acetobacterales | 35 | 0.0043 | 0.0325 |
| | KD4-96 | 1 | 0.0043 | 0.0325 |
| | 0319-7L14 | 1 | 0.0043 | 0.0325 |
| Semi-Arid | Cytophagales | 35 | 0.0025 | 0.0253 |
| | Isosphaerales | 35 | 0.0025 | 0.0253 |
| | uncultured | 0 | 0.0025 | 0.0253 |
| | Gaiellales | 1 | 0.0051 | 0.0303 |
| | KD4-96 | 1 | 0.0051 | 0.0303 |
| | Frankiales | 33 | 0.0101 | 0.0482 |
| | Rhizobiales | 32 | 0.0177 | 0.0482 |
| | Sphingomonadales | 32 | 0.0177 | 0.0482 |
| | Burkholderiales | 3 | 0.0177 | 0.0482 |

| | | | | |
|---|---|---|---|---|
| | Acetobacterales | 32 | 0.0177 | 0.0482 |
| | Vicinamibacterales | 3 | 0.0177 | 0.0482 |
| | Frankiales | 36 | 0.0022 | 0.0093 |
| | Cytophagales | 36 | 0.0022 | 0.0093 |
| | Gaiellales | 0 | 0.0022 | 0.0093 |
| | Chitinophagales | 36 | 0.0022 | 0.0093 |
| | uncultured | 0 | 0.0022 | 0.0093 |
| | IMCC26256 | 0 | 0.0022 | 0.0093 |
| | KD4-96 | 0 | 0.0022 | 0.0093 |
| | Rhodobacterales | 35 | 0.0043 | 0.0144 |
| Sub-Humid | 0319-7L14 | 1 | 0.0043 | 0.0144 |
| | Sphingomonadales | 34 | 0.0087 | 0.0216 |
| | Micromonosporales | 34 | 0.0087 | 0.0216 |
| | Vicinamibacterales | 2 | 0.0087 | 0.0216 |
| | Rhizobiales | 33 | 0.0152 | 0.0325 |
| | Micrococcales | 33 | 0.0152 | 0.0325 |
| | Solirubrobacterales | 4 | 0.0260 | 0.0458 |
| | Rubrobacterales | 4 | 0.0260 | 0.0458 |
| | Pyrinomonadales | 4 | 0.0260 | 0.0458 |

### Supplementary Methods

#### Method S1 - Soil analysis

Soil texture was determined by sedimentation analysis[5]. Specifically, 40 g of soil was added to 1 L of water at 25 °C in a 3 L cylindrical flask. The suspension was mixed thoroughly until homogenized and then allowed to settle. After 1 min, the height of the settled layer was recorded and taken as the sand fraction. To determine the silt fraction, the suspension was allowed to settle for 4 h and the increase in the settled layer height relative to the 1-min measurement was attributed to silt. Finally, after 9 h of settling, the height profile was measured again, and the additional increment was classified as the clay fraction.

#### Method S2 - Experimental design

Volatile organic compound (VOC) collection was performed under controlled environmental conditions using the flow-through cuvette system of Helmholtz Munich (Riedlmeier et al. 2017; Wenig et al. 2019). A schematic of the dynamic flow-through cuvette enclosure system can be found in Supplementary Fig. S3, and details of the measurement system can be found in (Riedlmeier et al. 2017), except that the original cuvettes were replaced by ten odorless polyethylene terephthalate (PET) bags (volume: 1 L) (Zhang et al. 2020) to allow simultaneous measurements during the rain simulation. To this end, a small section of the bag was covered with a 3 $cm^2$ adhesive patch to enable the glass syringe needle to pierce it. The (glass) syringe (Hamilton) was then used to introduce water to rewet the soil. Additionally, two more cuvettes were integrated into the system to allow continuous background monitoring and gas calibration.

Samples were enclosed under 12 h light/dark cycles (5-17 CET) at controlled temperatures of 28.4 ± 0.2 °C (max under light) and 22.8 ± 0.1 °C (min under dark), relative humidity of 11.2 ± 0.8% (light) and 65.2 ± 2.1% (dark), and light intensity of 135 ± 15 µmol photons $m^{-2}$ $s^{-1}$ photosynthetic photon flux density (PPFD). The diurnal changes of temperature due to the light cycles can be seen in Fig S4. Cuvette-enclosed samples were flushed continuously with purified air (200 ml $min^{-1}$; ZA30, LNI Swissgasat Srl, Italy), maintaining a constant and low total VOC concentration (<10 ppbv) and $CO_2$ levels at ~420 ppmv. From the cuvette outlet, the air stream was divided into three subsamples: 40 ml $min^{-1}$ was directed to a high-sensitivity proton-transfer-reaction quadrupole mass spectrometer (PTR-QMS) for real-time VOC analysis as previously described[6-9], 40 ml $min^{-1}$ to a thermo-desorption gas chromatography–mass spectrometry (TD-GC-MS) system for offline VOC analysis [6,7,9,10], and 80 ml $min^{-1}$ to an infrared gas analyzer (LI-850, Licor Biosciences GmbH, Hessen, Germany) for $CO_2$ and $H_2O$ measurements. Cuvettes were automatically switched every 6 minutes, resulting in a time resolution of 1 hour for real-time gas measurements. Of the 6-minute sampling interval, the first 4 minutes of data were discarded to eliminate potential carryover from the preceding cuvette, and the remaining 2 minutes were background corrected using data from blanks (see below) and averaged for analysis of volatile fluxes.

The experiments were designed to monitor VOC emissions and gas exchange before and after two simulated rain events, the first after a long drought period (>3 months) and the second after a short (31 days) desiccation period during which the soil water content (SWC) returned to its pre-rain condition. The protocol proceeded as follows: on the day before the first rain simulation (day 0), samples were enclosed between 09:30 and 10:00 CET for baseline (dry soil) measurements. GC-MS sampling was conducted for 17 hours from 17:00 on day 0 to 10:00 the following day (day 1). On day 1, between 16:00 and 16:30, 25 ml of Milli-Q water was applied slowly to the soil surface using a glass syringe (Hamilton), in five sequential injections of 5 ml each at a rate of 1.5 ml $min^{-1}$. The syringe needle pierced the cuvette wall, and the same procedure was performed on blank cuvettes to subtract the background. After wetting, GC-MS sampling started from 17:00 on day 1 and continued until 10:00 on day 2. The experimental timeline is depicted in Fig. S5.

Of the five replicate samples, VOC emissions and gas exchange were monitored online for three replicates through day 4. Thereafter, samples were removed from the cuvettes and allowed to dry for 31 days to achieve a similar SWC to the initial values as initially, after which the entire measurement protocol was repeated following a second rain event. All samples were measured in a fully randomized order to avoid potential cuvette effects. Soil samples were photographed at the end of the experiments (Suppl. Fig S6).

**Method S3 - VOC analysis**

Volatile organic compound (VOC) emissions were measured simultaneously using a high-sensitivity proton-transfer-reaction quadrupole mass spectrometer (PTR-QMS; Ionicon Analytik GmbH) and thermal desorption gas chromatography – mass spectrometry (TD-GC–MS, Gerstel; GC, 7890A and MS, 5975C both from Agilent Technologies), following previously established protocols [8,9,11] detailed below.

**PTR-QMS analysis**

PTR-QMS was operated at an E/N of 135 Td (E = the electric field strength, N = the gas number density; 1 Td = $10^{-17}$ V $cm^2$) with a drift tube (dt) pressure of 2.2 mbar; dt voltage of 600 V, dt temperature of 60°C. The relative abundance of $H_3O^+(H_2O)_1$, $O_2^+$, and $NO^+$ ions were maintained below 10%, 3%, and 0.2% of the primary ions, respectively. Calibration was conducted using humidity-dependent dilution (0-90% RH at 24°C) across 12 concentrations (0-150 ppbv) of an 11-VOC standard mixture (Apel-Riemer Environmental), passed through the entire analytical system as described before [7].

A list of the monitored protonated masses and their chemical assignments based on TD-GC-MS data is provided in Suppl. Table S2. The ion at m/z 81 $[C_6H_9]^+$, which can originate from a range of compounds including monoterpenes, sesquiterpenes, geosmin, and various unsaturated C6 alcohols or aldehydes, did not directly correlate to a specific VOC class; it is therefore presented as representative of $[C_6H_9]^+$ and has multiple potential VOC sources. Sesquiterpenes were quantified using empirically derived transmission factors[12] calculated from instrumental sensitivities based on a 17-VOC standard mixture (Ionicon). Results showed excellent agreement (<5%) with TD-GC–MS calibrations [9]. Calibration and gas standard uncertainties were estimated at <10%. Limits of detection (LOD), defined as 2σ, ranged between 0.13 and 5.4 ppbv.

#### TD-GC-MS analysis

VOC chemical characterization was performed using TD-GC–MS. Air samples were collected in glass cartridges filled with Tenax TA and Carbopack X [13], spiked with δ-2-carene as an internal standard [9]. The method followed prior studies [14] [11] [8].

Cartridges were flushed with ultrapure helium (2.5 min, 100 mL $min^{-1}$, 25°C) and desorbed at 250°C (280°C $min^{-1}$ ramp, held for 5 min). Desorbed VOCs were refocused in a cryo-injection system (CIS) at -50°C (100 mL $min^{-1}$, 15 psi vent pressure), then thermally desorbed by ramping to 270°C at 12°C $s^{-1}$ and held for 4 min in splitless mode.

Chromatographic separation was achieved using a DB-5MS column (60 m × 250 µm × 0.25 µm + 10 m DG; Agilent Technologies), with helium (purity 5.0) as the carrier gas at 1 mL $min^{-1}$. The GC oven temperature was programmed as follows: 40°C (start), ramped to 150°C at 10°C $min^{-1}$, to 175°C at 80°C $min^{-1}$, then to 190°C and subsequently 250°C at 80°C $min^{-1}$, and finally to 300°C at 100°C $min^{-1}$ (held for 6 min). Ionization was by electron impact (70 eV) and detection by quadrupole mass spectrometry. The MS source and quadrupole temperatures were 230°C and 150°C, respectively. Data were acquired in both scan mode (m/z 35-250) and single-ion mode (SIM), targeting ions at m/z 93, 121, 136, 161, and 204 (dwell time: 25-50 ms).

Untargeted data analysis was conducted using MS-DIAL [15]. Data were converted to AIA (.CDF) and Abf formats, then processed for peak detection, alignment, filtering, and deconvolution. Final identification and quantification were performed manually, using pure standards, library matches (NIST 20 and Wiley 275 GC-MS), and calibration curves. Kovats retention indices were calculated using a $C_9$-$C_{25}$ alkane standard (Sigma-Aldrich, Germany).

Quantification relied on calibration curves prepared from six concentrations of pure standard mixtures (monoterpenoids, sesquiterpenoids, benzenoids, alkanes, ketones), each prepared in triplicate [9]. Separate calibrations were conducted for geosmin and 2-MIB. Signal linearity was excellent ($r^2$ = 0.986-0.9993) across the measured concentration range. Instrument sensitivity drift was corrected using the δ-2-carene internal standard. Blank replicates were used for background correction. Detection and quantification limits were set at 2σ and 3×LOD, respectively [13].

#### Method S4 - Statistical analysis of bacterial diversity and community composition

All statistical analyses were performed on the 16S rRNA amplicon dataset after restricting the ASV table to bacteria based on the taxonomic assignment. Taxa with zero counts across all samples were removed. For multivariate analyses, ASV counts were converted to relative abundances per sample and then Hellinger-transformed (square root of relative abundance) to reduce the influence of very rare taxa and to make the data more suitable for Euclidean-based methods[16].

To assess differences in community composition, we used a PERMANOVA framework on the Hellinger-transformed ASV matrix[17]. Euclidean distances among samples in Hellinger space were used as the dissimilarity measure, which is equivalent to performing PERMANOVA on Hellinger-transformed community data. We tested the effects of habitat type (biocrust vs. topsoil) and climatic region (sub-humid, semi-arid, arid, hyper-arid) on community composition using permutation-based multivariate analysis of variance with 999 permutations. The proportion of variance explained by each factor was expressed as $R^2$. Because PERMANOVA can be sensitive to heterogeneity of dispersion, we also quantified within-group dispersion in the same Hellinger space (PERMDISP [18]). For each factor (*Type* and *Region*), we calculated the distance of each sample to its group centroid and compared mean distances among groups using an ANOVA-like test with 999 permutations (PERMDISP analogue). This allowed us to evaluate whether observed PERMANOVA effects reflected differences in group centroids (location) rather than solely differences in within-group variability (dispersion).

To relate these multivariate patterns to the order-level relative abundance profiles (Fig. 1), we aggregated the bacterial ASV counts to the order level and recalculated relative abundances per sample. We then identified 'dominant' orders as those with a mean relative abundance of ≥ 1% across all samples; only these orders were retained for univariate tests associated with the stacked-bar figure. For each dominant order, we tested for effects of climatic region within each habitat type using Kruskal-Wallis tests [18], and for effects of habitat type within each climatic region using Wilcoxon rank-sum (Mann-Whitney U) tests[19]. To control for multiple comparisons, p-values were adjusted using the Benjamini–Hochberg false discovery rate (FDR) procedure within each factor[20]. Orders with FDR-corrected $q < 0.05$ were considered to respond significantly to aridity and/or habitat. All statistical computations were carried out in a scripted environment (R or Python; vegan-like PERMANOVA/PERMDISP and non-parametric tests as described above), ensuring that the same abundance and metadata tables used to generate the relative-abundance figure were used consistently for the multivariate and univariate analyses.

3 USDA-NRCS. Java Newhall Simulation Model (jNSM) 1.6.0 User Guide – Part I. Natural Resources Conservation Service. jNSM 1.6.0 User Guide – Part II: Useful References. Natural Resources Conservation Service. . *hxps://www.nrcs.usda.gov/resources/educaVon-and-teaching-materials/java-newhallsimulaVon-model-jnsm*

4 Bay, S. K. *et al.* Soil Bacterial Communities Exhibit Strong Biogeographic Patterns at Fine Taxonomic Resolution. *mSystems* **5**, doi:10.1128/mSystems.00540-20 (2020).

5 Smith, K. A., Mullins, C. E. *Soil Analysis: Physical Methods*. (1991).

6 Ghirardo, A. *et al.* Determination of de novo and pool emissions of terpenes from four common boreal/alpine trees by 13CO2 labelling and PTR-MS analysis. *Plant Cell Environ* **33**, 781-792, doi:10.1111/j.1365-3040.2009.02104.x (2010).

7 Ghirardo, A., Gutknecht, J., Zimmer, I., Bruggemann, N. & Schnitzler, J. P. Biogenic volatile organic compound and respiratory CO2 emissions after 13C-labeling: online tracing of C translocation dynamics in poplar plants. *PLoS One* **6**, e17393, doi:10.1371/journal.pone.0017393 (2011).

8 Ghirardo, A., Heller, W., Fladung, M., Schnitzler, J. P. & Schroeder, H. Function of defensive volatiles in pedunculate oak (Quercus robur) is tricked by the moth Tortrix viridana. *Plant Cell Environ* **35**, 2192-2207, doi:10.1111/j.1365-3040.2012.02545.x (2012).

9 Ghirardo, A. *et al.* Origin of volatile organic compound emissions from subarctic tundra under global warming. *Glob Chang Biol* **26**, 1908-1925, doi:10.1111/gcb.14935 (2020).

10 Kreuzwieser, J. *et al.* The Venus flytrap attracts insects by the release of volatile organic compounds. *J Exp Bot* **65**, 755-766, doi:10.1093/jxb/ert455 (2014).

11 Ghirardo, A. *et al.* Urban stress-induced biogenic VOC emissions and SOA-forming potentials in Beijing. *Atmos Chem Phys* **16**, 2901-2920, doi:10.5194/acp-16-2901-2016 (2016).

12 Taipale, R. *et al.* Technical Note: Quantitative long-term measurements of VOC concentrations by PTR-MS – measurement, calibration, and volume mixing ratio calculation methods. *Atmos Chem Phys* **8**, 6681-6698, doi:10.5194/acp-8-6681-2008 (2008).

13 Ghirardo, A. & Brilli, F. in *Monitoring Forest Damage with Metabolomics Methods* 127-186 (2024).

14 Weikl, F., Ghirardo, A., Schnitzler, J. P. & Pritsch, K. Sesquiterpene emissions from Alternaria alternata and Fusarium oxysporum: Effects of age, nutrient availability, and co-cultivation. *Sci Rep* **6**, 22152, doi:10.1038/srep22152 (2016).

15 Tsugawa, H. *et al.* MS-DIAL: data-independent MS/MS deconvolution for comprehensive metabolome analysis. *Nat Methods* **12**, 523-526, doi:10.1038/nmeth.3393 (2015).

16 Legendre, P. & Gallagher, E. D. Ecologically meaningful transformations for ordination of species data. *Oecologia* **129**, 271-280, doi:10.1007/s004420100716 (2001).

17 Anderson, M. J. A new method for non-parametric multivariate analysis of variance. *Austral Ecology* **26**, 32-46, doi:https://doi.org/10.1111/j.1442-9993.2001.01070.pp.x (2001).

18 Anderson, M. J. Distance-Based Tests for Homogeneity of Multivariate Dispersions. *Biometrics* **62**, 245-253, doi:10.1111/j.1541-0420.2005.00440.x (2005).

19 Mann, H. B. & Whitney, D. R. On a test of whether one of two random variables is stochastically larger than the other. *Annals of Mathematical Statistics* **18**, 50-60, doi:10.1214/aoms/1177730491 (1947).
20 Benjamini, Y. & Hochberg, Y. Controlling the False Discovery Rate: A Practical and Powerful Approach to Multiple Testing. *Journal of the Royal Statistical Society: Series B (Methodological)* **57**, 289-300, doi:https://doi.org/10.1111/j.2517-6161.1995.tb02031.x (1995).
